# Bidirectional substrate shuttling between the 26S proteasome and the Cdc48 ATPase promotes protein degradation

**DOI:** 10.1101/2023.12.20.572403

**Authors:** Hao Li, Zhejian Ji, Joao A. Paulo, Steven P. Gygi, Tom A. Rapoport

## Abstract

Most eukaryotic proteins are degraded by the 26S proteasome after modification with a polyubiquitin chain. Substrates lacking unstructured segments cannot be degraded directly and require prior unfolding by the Cdc48 ATPase (p97 or VCP in mammals) in complex with its ubiquitin-binding partner Ufd1-Npl4 (UN). Here, we use purified yeast components to reconstitute Cdc48-dependent degradation of well-folded model substrates by the proteasome. We show that a minimal system consists of the 26S proteasome, the Cdc48-UN ATPase complex, the proteasome cofactor Rad23, and the Cdc48 cofactors Ubx5 and Shp1. Rad23 and Ubx5 stimulate polyubiquitin binding to the 26S proteasome and the Cdc48-UN complex, respectively, allowing these machines to compete for substrates before and after their unfolding. Shp1 stimulates protein unfolding by the Cdc48-UN complex, rather than substrate recruitment. *In vivo* experiments confirm that many proteins undergo bidirectional substrate shuttling between the 26S proteasome and Cdc48 ATPase before being degraded.

## INTRODUCTION

In eukaryotes, the majority of regulated protein degradation is carried out by the 26S proteasome, and substrate proteins are typically first modified by the attachment of a polyubiquitin chain (Collins and Goldberg, 2017). In many cases, the ubiquitin chain is linked to a misfolded protein, which can then be degraded directly by the proteasome. However, in other cases, the substrate is well-folded or present in membranes or complexes, and it therefore requires initial unfolding by an AAA ATPase, called Cdc48 in yeast and p97 or VCP in mammals. Examples include endoplasmic reticulum (ER)–associated protein degradation (ERAD) (Bodnar and Rapoport, 2017a), the extraction of actin and myosin from myofibrils during muscle atrophy (Piccirillo and Goldberg, 2012), and the removal of various proteins from chromatin (van den Boom and Meyer, 2018). p97 has also been implicated in the proteasomal degradation of amyloid-like aggregates of the microtubule-associated protein tau (Saha et al., 2022), but a systematic search for Cdc48/p97-dependent proteasomal substrates has not yet been performed. Consistent with a central role in protein quality control, mutations in human p97 cause degenerative diseases in multiple organs and tissues (Meyer and Weihl, 2014; Tang and Xia, 2016).

The 26S proteasome consists of a 19S regulatory particle and a 20S proteolytic core (Collins and Goldberg, 2017). The 19S particle first binds the polyubiquitin chain attached to a substrate (**figure S1A**). Next, a flexible segment of the substrate inserts into the central pore of the hexameric ATPase ring at the base of the 19S particle (Davis et al., 2021); without such a flexible segment, a substrate cannot be efficiently degraded. The ubiquitin chain is then removed by the deubiquitinase (DUB) Rpn11 (Verma et al., 2002), and the polypeptide is pushed by the ATPases into the proteolytic chamber of the 20S particle.

Cdc48/p97 consists of an N-terminal (N) domain, and two ATPase domains called D1 and D2. Six Cdc48 molecules assemble into a double-ring structure with a central pore (**figure S1B**) (Bodnar and Rapoport, 2017a; DeLaBarre and Brunger, 2003). Cdc48/p97 cooperates with a number of conserved cofactors, in particular the heterodimeric Ufd1-Npl4 (UN) complex, which functions in ERAD and many other processes (Ye et al., 2017). A single UN dimer associates with the hexameric ATPase and recruits substrates through attached Lys48-linked ubiquitin chains (Bruderer et al., 2004). Substrate processing by the Cdc48-UN complex is initiated by the ATP-independent unfolding of one of the ubiquitin molecules in the chain (i.e., the initiator ubiquitin) (Twomey et al., 2019). The N-terminus of the initiator ubiquitin inserts into the central pore across both hexameric ATPase rings (**figure S1B**). The ATPase then uses the energy of ATP hydrolysis to sequentially move the initiator ubiquitin, all ubiquitin molecules positioned between the initiator and the substrate, and finally the substrate itself through the pore, causing their unfolding (Ji et al., 2022). Following translocation, ubiquitin molecules refold, whereas the substrate generally remains unfolded, although in some cases, it can spontaneously refold. The polyubiquitin chain allows a substrate to rebind to the Cdc48-UN complex, so that the ATPase can act again on a substrate that has already been translocated (Ji *et al*., 2022).

It is generally assumed that the Cdc48/p97 ATPase acts upstream of the 26S proteasome (Tsuchiya et al., 2020); a folded protein would be unfolded by Cdc48/p97 and then be transferred by shuttling factors to the proteasome for degradation (Finley, 2009). In *S. cerevisiae*, three conserved proteins have been proposed to function as shuttling factors: Rad23, Dsk2, and Ddi1 (Tsuchiya *et al*., 2020). All three proteins contain ubiquitin-binding UBA domains as well as proteasome-binding UBL domains (UBA-UBL proteins), and can therefore recruit polyubiquitinated substrates to the proteasome. Rad23 and Dsk2 stimulate the degradation of ubiquitinated substrates (Elsasser et al., 2004; Lambertson et al., 1999; Medicherla et al., 2004; Rao and Sastry, 2002; Richly et al., 2005; Shi et al., 2016; Verma et al., 2004), but the mechanism by which the UBA-UBL proteins participate in Cdc48-dependent protein degradation remains poorly understood. For example, the UBA-UBL proteins might actually inhibit the degradation of folded substrates, as they could sequester these proteins on the proteasome and prevent their unfolding by the Cdc48 ATPase. Conversely, the Cdc48/p97 ATPase might interfere with the degradation of misfolded proteins, as these might bind through their ubiquitin chain to the Cdc48/p97-UN complex, where they would undergo cycles of unfolding and become sequestered from degradation by the proteasome. Thus, it remains unclear how substrates would avoid being trapped on either the proteasome or the Cdc48/p97 ATPase.

A mechanistic understanding of Cdc48 ATPase-dependent protein degradation is difficult to achieve by experiments with intact cells or crude extracts alone, as such systems contain too many interacting and interfering components. *In vitro* systems are needed, in which the regulated degradation of proteins can be reproduced with a minimal set of purified components. Once such components have been identified, they can be tested in a more direct way by *in vivo* experiments. An initial attempt to recapitulate *in vitro* Cdc48-dependent proteasomal degradation of a folded protein resulted in low efficiency (<10% degradation over 30 min) and did not explore the role of cofactors (Olszewski et al., 2019).

Here, we report the establishment of an efficient reconstituted system that recapitulates the degradation of well-folded proteins. Our results lead to a novel model for protein degradation in which the 26S proteasome and Cdc48 ATPase utilize cofactors to compete for polyubiquitinated substrates before and after their unfolding. *In vivo* experiments confirm that the two molecular machines bidirectionally shuttle substrates between them, ultimately ensuring that folded substrates are first unfolded and then degraded.

## RESULTS

### Experimental strategy

The goal of this study is to identify the minimum components required for the degradation of well-folded proteins. We start out with the simplest reconstituted system that consists of only the 26S proteasome and the Cdc48 ATPase complex and then test candidate proteins that may participate in the interplay of the two molecular machines. All components were from *S. cerevisiae* (for purity, see **figures S1C; S1D**). To mimic the situation in intact cells, we used them at concentrations approximating those in *Saccharomyces cerevisiae* cells (∼250-500 nM), estimates that are based on quantitative proteomics data (**figure S1E**) and a volume of 42 μm^3^ for a yeast cell (Jorgensen et al., 2002). *In vivo* experiments were subsequently performed to test whether the conclusions drawn from the *in vitro* reconstitutions apply to *S. cerevisiae* cells.

### Cdc48-dependent degradation of a folded substrate by the proteasome

We first identified a substrate with a flexible segment, which could be directly degraded by purified 26S proteasomes from *S. cerevisiae* (Lander et al., 2012) without addition of any other component. The substrate consists of a disordered polypeptide segment of 43 amino acids, fused to the N terminus of superfolder green fluorescent protein (sfGFP) and a C-terminal flexible segment of 50 amino acids (Ji *et al*., 2022) (**Figure 1A**). Both the N- and C-terminal unstructured segments are long enough to insert into the 19S particle of the proteasome and initiate translocation into the proteolytic chamber (Inobe et al., 2011). To facilitate detection of degradation products, the fusion protein was labeled with the fluorescent dye Dylight800 on cysteines. A single chain of Lys48-linked ubiquitin molecules was attached to a lysine residue within the N-terminal flexible segment (Ji *et al*., 2022), and the modified protein was purified by gel filtration to enrich for chains of 10-25 ubiquitins. The substrate is referred to as Ub(n)-TAIL (**Figure 1A**). Incubation of Ub(n)-TAIL with purified 26S proteasomes, ATP, and an ATP-regenerating system caused its efficient degradation (**Figure 1B**). The decrease of fluorescence in the precursor quantitatively matched the appearance of fluorescence in peptides, indicating that the substrate was degraded and not just deubiquitinated. The reaction was dependent on the presence of ATP (**Figure 1B**) and essentially complete within 20 min (**figure S2A)**. Degradation was inhibited by the addition of o-phenanthroline (**Figure 1B**, lane 4), which targets the proteasomal DUB Rpn11 (Verma *et al*., 2002), consistent with removal of the ubiquitin chain being required for polypeptide translocation into the proteolytic chamber (Bard et al., 2019). No degradation was observed with the 20S core particle (**figure S2B**). These data demonstrate that the purified 26S proteasomes can efficiently degrade a polyubiquitinated protein containing a disordered segment. Two other tested substrates with flexible segments at either the N- or C-terminus (Ub(n)-NTAIL and Ub(n)-CTAIL) were not efficiently degraded (**figures S2C,D and S2E,F**), perhaps because the distance between the ubiquitin chain and the flexible segment or other features of the substrate were suboptimal. Thus, consistent with results in the literature (Davis et al., 2023; Martinez-Fonts et al., 2020), not all polyubiquitinated proteins with flexible segments are efficiently degraded by the 26S proteasome alone.

**Figure 1.**
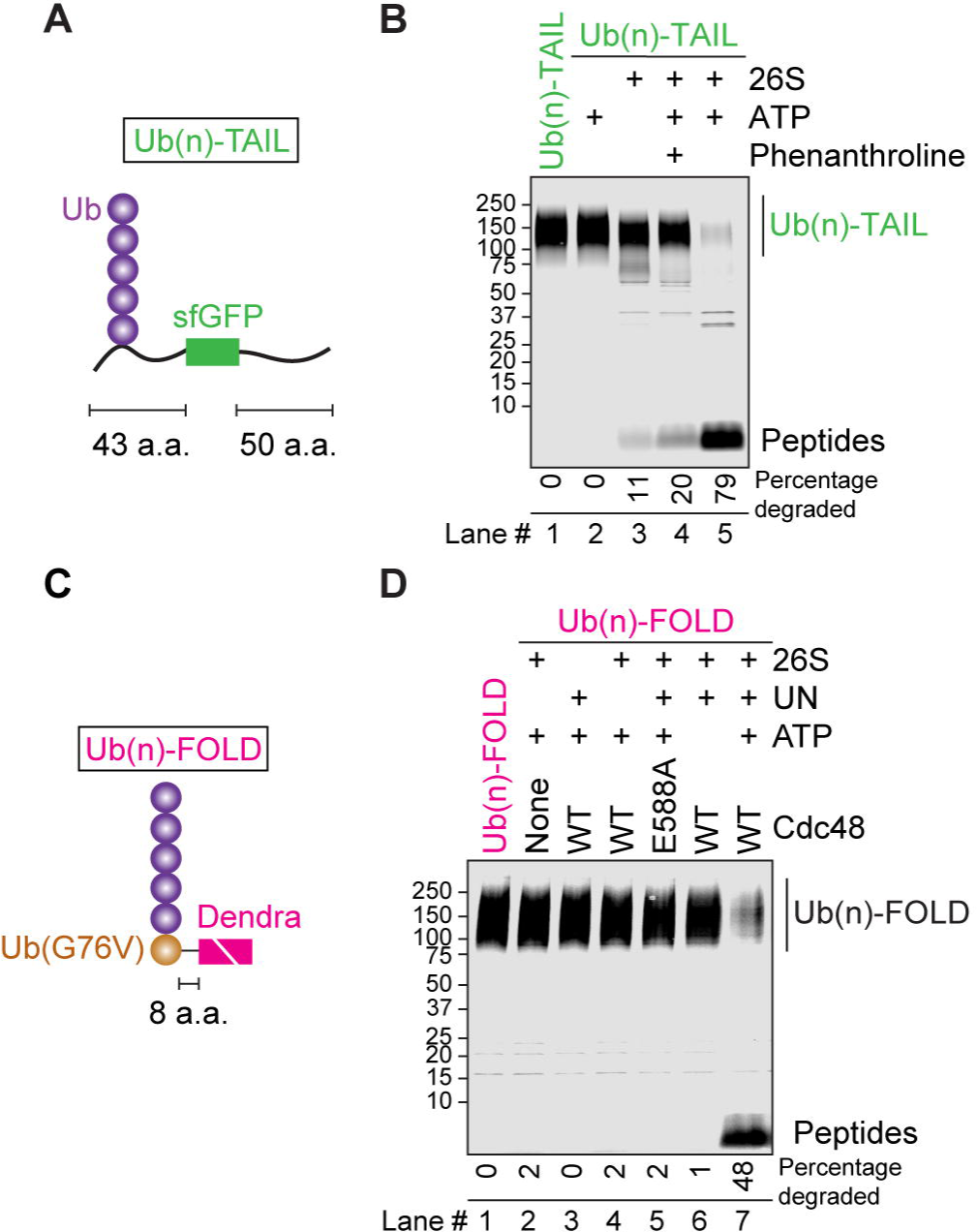
Cdc48-dependent degradation of a folded model substrate. (**A**) Scheme of the model substrate Ub(n)-TAIL, which contains superfolder GFP (sfGFP), flexible tails at the N- and C-terminus, and a single chain of K48-linked ubiquitin (Ub) molecules. (**B**) Degradation of Ub(n)-TAIL, labeled with the fluorescent dye Dylight 800, by 26S proteasomes. Where indicated, ATP or o-phenanthroline was added. Reactions were incubated for 60 min and analyzed by SDS-PAGE and fluorescence scanning. The percentage of substrate degraded was quantified by determining the fluorescence in peptides (after background subtraction) and comparing it to the total fluorescence in each lane of the SDS gel. The total fluorescence in the lanes was the same within a range of +/− 15%. (**C**) Scheme of the well-folded model substrate Ub(n)-FOLD. The ubiquitin (Ub) mutant Ub-G76V was fused to Dendra with an 8 amino acid linker. Dendra was photo-cleaved into two polypeptide fragments (indicated by a white line). A ubiquitin chain was attached to Ub-G76V. (**D**) Degradation of Dylight 800-labeled, photo-cleaved Ub(n)-FOLD by 26S proteasomes in the presence of Cdc48-UN complex. Cdc48 was either wild-type (WT) or contained a mutation (E588A) that abolished its unfolding activity. Quantification was done as in (A). The total fluorescence in the lanes was the same within a range of +/− 10%.

Next, we designed a well-folded substrate that must first be unfolded by the Cdc48 ATPase before it can be degraded by the 26S proteasome. To mimic the behavior of many endogenous substrates, we first chose a protein that is irreversibly unfolded by Cdc48. The substrate was generated by fusing ubiquitin to the fluorescent protein Dendra with a linker of 8 amino acids that is too short to initiate processing by the 26S proteasome (Inobe *et al*., 2011). The last amino acid of ubiquitin was mutated to valine (i.e., G76V) to preclude the separation of the fusion partners by Rpn11. The substrate was subjected to UV irradiation, which splits the Dendra moiety into two fragments that are irreversibly separated after unfolding by Cdc48 (Ji *et al*., 2022; Kaberniuk et al., 2017). A cysteine in the linker was labeled with Dylight800, and a single Lys48-linked ubiquitin chain was attached to the N-terminal ubiquitin (Ji *et al*., 2022). The resulting polyubiquitinated substrate was enriched by gel filtration for chains of 10-25 ubiquitin molecules, and is referred to as Ub(n)-FOLD (**Figure 1C**).

As expected, Ub(n)-FOLD was only degraded when the Cdc48 ATPase and its UN cofactor were added to the reaction containing 26S proteasomes (**Figure 1D**, lane 7 versus 2). The majority of the substrate was degraded within 30 min (**figure S2G**). Again, the decrease of fluorescence in the precursor quantitatively matched the appearance of fluorescence in peptides. No degradation was observed when Cdc48, UN, or ATP were omitted (**Figure 1D**, lanes 3, 4, 6). Importantly, degradation was blocked when the unfolding activity of Cdc48 was inactivated by a mutation in the D2 ATPase domain (Bodnar and Rapoport, 2017a) (E588A; lane 5). Thus, the well-folded substrate must first be unfolded by the Cdc48 ATPase before it can be degraded by the proteasome. The unfolding activity of the ATPase complex was confirmed by incubating Cdc48-UN with Ub(n)-FOLD in the absence of proteasomes; a gradual decrease of fluorescence was observed (**figure S2H**), caused by the irreversible separation of the fragments of photo-cleaved Dendra (Bodnar and Rapoport, 2017a). Taken together, these data show that the efficient degradation of a polyubiquitinated, folded substrate can be recapitulated with a system consisting of only the Cdc48-UN ATPase complex and the 26S proteasome.

### Effect of the proteasome cofactor Rad23 on Cdc48-dependent protein degradation

Next we examined the effect of the UBA-UBL shuttling factors Rad23, Dsk2, and Ddi1, which are thought to mediate the transfer of unfolded substrates from Cdc48 to the proteasome (Tsuchiya *et al*., 2020). We first used pull-down experiments to test whether these factors recruit substrates to the proteasome. Because Ddi1 is also a protease that can cleave Ub(n)-TAIL (Yip et al., 2020), the experiments with this substrate employed a proteolytically inactive Ddi1 mutant (Ddi1 D220N). Rad23 was found to be a potent substrate recruitment factor (**Figure 2A**, lane 7). The binding of Rad23 and substrate to the proteasome was synergistic, as Rad23 strongly potentiated the recruitment of Ub(n)-FOLD and Ub(n)-TAIL (**Figure 2A**, lane 7 versus 3; **figure S3A**, lane 7 versus 3), and *vice versa*, substrate stimulated the binding of Rad23 (**Figure 2A** and **figure S3A,** anti-SBP immunoblots). Surprisingly, Dsk2 did not strongly promote substrate recruitment to the proteasome (**Figure 2A** and **figure S3A,** lanes 8), even though it can do so *in vivo* (Tsuchiya et al., 2017). Ddi1 was even slightly inhibitory (**Figure 2A** and **figure S3A,** lanes 9). As expected from the presence of ubiquitin-binding UBA domains, all three factors bound similarly well to polyubiquitinated substrate in the absence of proteasomes (**figure S3B**).

**Figure 2.**
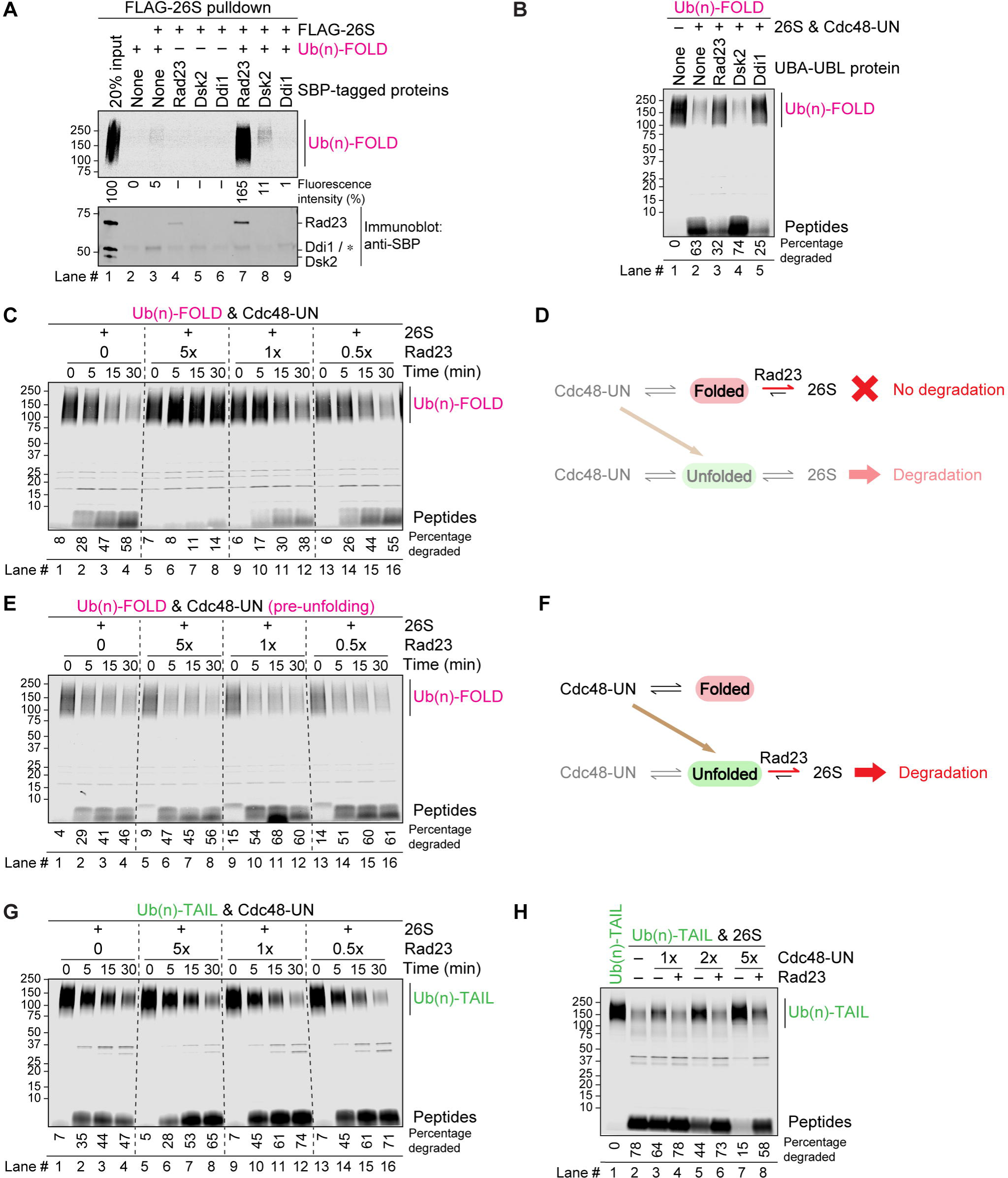
Effect of Rad23 on the degradation of model substrates. (**A**) Rad23, Dsk2, and Ddi1 were tested for substrate recruitment to the 26S proteasome. FLAG-tagged 26S proteasomes were incubated with fluorescently labeled Ub(n)-TAIL and SBP-tagged versions of Rad23, Dsk2, or Ddi1, as indicated. After immunoprecipitation with FLAG antibodies, the samples were analyzed by SDS-PAGE followed by fluorescence scanning and blotting for SBP. The numbers below the gel give the intensity of the fluorescent bands relative to 20% input. (**B**) Fluorescently labeled, photo-converted Ub(n)-FOLD was incubated with 26S proteasomes, Cdc48-UN, and the UBA-UBL proteins Rad23, Dsk2, or Ddi1, as indicated. After 60 min, the samples were analyzed by SDS-PAGE and fluorescence scanning. The percentage of substrate degraded was quantified by determining the fluorescence in peptides (after background subtraction) and comparing it to the total fluorescence in each lane of the SDS gel. (**C**) As in (B), but time course of degradation with different Rad23 concentrations, given relative to the concentrations of proteasomes and Cdc48-UN. (**D**) Scheme explaining the inhibitory effect of Rad23 on the degradation of Ub(n)-FOLD. Rad23 sequesters the substrate on the 26S proteasome, so that it can neither be degraded nor be transferred to Cdc48-UN for unfolding. (**E**) Photo-converted Ub(n)-FOLD was first unfolded by Cdc48-UN (**figure S3D**). Proteasomes and Rad23 were then added and substrate degradation assessed as in (C). Note the degradation in the presence of even high concentrations of Rad23 (compare lanes 5-8 in panels (E) and (C)). (**F**) Scheme explaining the lack of Rad23 inhibition in (E). Recruitment of the unfolded protein to the proteasome allows degradation. (**G**) As in (C), but with the direct proteasome substrate Ub(n)-TAIL. Note that, as in (E), Rad23 does not inhibit substrate degradation. (**H**) Proteasomal degradation of Ub(n)-TAIL in the presence of stoichiometric concentrations of Rad23 and increasing concentrations of Cdc48-UN complex. 1x refers to stoichiometric concentrations with respect to 26S proteasomes. Quantification was done as in (B).

Because Rad23 was most potent in substrate recruitment to the 26S proteasome, we asked whether it would stimulate the degradation of the folded model substrate, Ub(n)-FOLD, when added together with 26S proteasomes and the Cdc48-UN complex. Unexpectedly, Rad23 was inhibitory (**Figure 2B**, lane 3 versus 2), and almost completely blocked degradation when added in excess over Cdc48-UN and 26S proteasomes (**Figure 2C**). Rad23 also reduced the rate of substrate unfolding by Cdc48-UN in the absence of proteasomes (**figure S3C**), in part because it can sequester polyubiquitinated substrates (**figure S3B**). In the context of the proteasome, however, Rad23 likely inhibited degradation mainly by promoting the formation of a non-productive complex between the 26S proteasome and the folded protein, such that the substrate could neither be degraded nor be transferred to Cdc48-UN for unfolding (**Figure 2D**). This model is supported by experiments in which we pre-incubated Ub(n)-FOLD with the Cdc48-UN complex to unfold the substrate (**figure S3D**), and then added Rad23 and 26S proteasomes. In this case, no inhibition of substrate degradation was observed, even with a large excess of Rad23 (**Figure 2E**). Rad23 even slightly stimulated degradation, likely by recruiting the unfolded substrate to the proteasome (**Figure 2F**). Indeed, when Cdc48-mediated unfolding was bypassed by using the Ub(n)-TAIL substrate, Rad23 showed no inhibition and instead moderately potentiated degradation (**Figure 2G**). Rad23 also had a negligible effect on the degradation of Ub(n)-TAIL in the absence of Cdc48-UN **(figure S3E**, lane 3 versus 2). These results thus indicate that Rad23 can recruit both folded and unfolded polyubiquitinated proteins to the proteasome (**Figures 2D; 2F**). They suggest that the Cdc48 complex does not obligatorily function upstream of the proteasome, as hitherto assumed.

Our model predicts that excessive substrate recruitment to the Cdc48 complex would also suppress degradation, as it would sequester substrate away from the proteasome. Indeed, proteasomal degradation of the Ub(n)-TAIL substrate was progressively inhibited by increasing the concentration of Cdc48-UN (**Figure 2H**; lanes 2, 3, 5, 7). Importantly, Rad23 counteracted the inhibition exerted by excess Cdc48-UN (**Figure 2H**; lanes 4, 6, 8). Taken together, our results demonstrate that the 26S proteasome and Cdc48-UN complex compete for polyubiquitinated substrates, with Rad23 shifting the balance towards the proteasome.

### A role for the Cdc48-cofactor Ubx5 in protein degradation

We reasoned that there must be a factor that counteracts the effect of Rad23 and shifts the balance in the opposite direction, i.e., towards the Cdc48 ATPase complex. The best candidate for the missing component would be one of the UBA-UBX proteins Shp1, Ubx2, and Ubx5, which contain both a ubiquitin-binding UBA domain and a Cdc48-interacting UBX domain (Schuberth and Buchberger, 2008).

We first asked whether any of these factors could help recruit substrates to the Cdc48-UN complex. All three UBA-UBX proteins interacted similarly well with polyubiquitinated substrate in the absence of Cdc48 (**figure S3B**), as expected from their UBA domains. In the presence of Cdc48-UN, we found that Ubx5, and to a lesser extent Ubx2, potentiated substrate recruitment to Cdc48-UN (**figure S3F**, lanes 8 and 9 versus 3), whereas Shp1 did not (lane 7). Notably, Ubx5 and substrate bound synergistically to Cdc48-UN, as one component enhanced the binding of the other (**figure S3F**; lane 9 versus 3 and 6), analogously to the synergistic binding of Rad23 and substrate to the proteasome (**Figure 2A**).

Ubx5 inhibited the degradation of Ub(n)-TAIL in a concentration-dependent manner (**Figure 3A**, lanes 2-5), but had only a small effect in the absence of Cdc48-UN (**figure S3E**, lane 4 versus 2). Inhibition was also observed with Ub(n)-FOLD (**Figure 3B**; lanes 2-5), even though unfolding was not affected (**figure S3G**). Thus, the addition of Ubx5 promotes the binding of both folded and unfolded substrates to the Cdc48-UN complex and prevents substrate transfer from Cdc48-UN to the 26S proteasome (**Figure 3C**).

**Figure 3.**
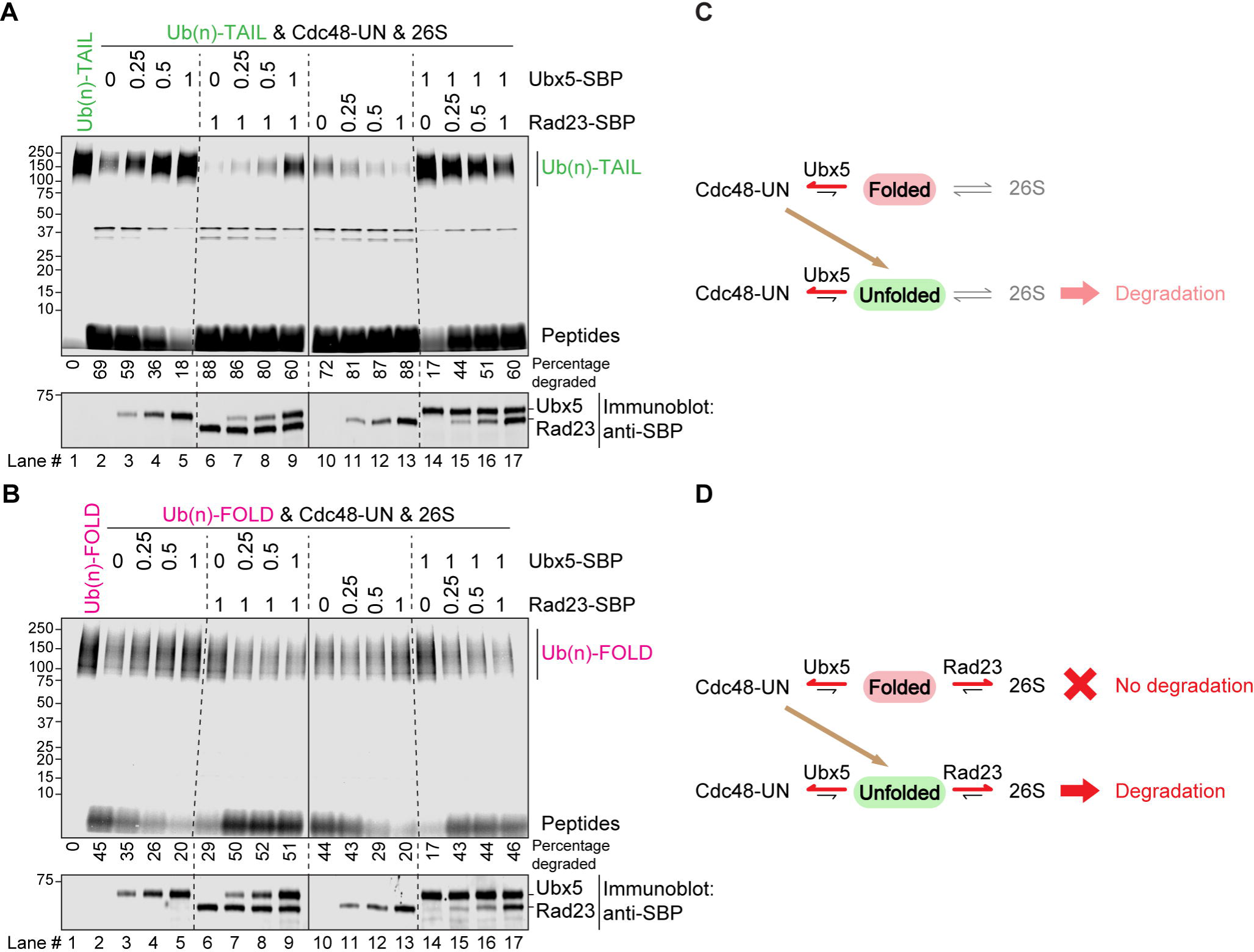
Ubx5 and Rad23 counteract one another in protein degradation. (**A**) Fluorescently labeled Ub(n)-TAIL was incubated with 26S proteasomes, Cdc48-UN, and different concentrations of SBP-tagged Ubx5 or SBP-tagged Rad23. The concentrations of the cofactors are given relative to that of 26S proteasomes. After 60 min, the samples were analyzed by SDS-PAGE, followed by fluorescence scanning and blotting for SBP. The percentage of substrate degraded was quantified by determining the fluorescence in peptides (after background subtraction) and comparing it to the total fluorescence in each lane of the SDS gel. (**B**) As in (A), but with photo-converted Ub(n)-FOLD substrate. (**C**) Scheme explaining the inhibition observed in (A) and (C) when only Ubx5 was added. Ubx5 sequesters folded and unfolded substrates on Cdc48-UN, preventing their transfer to the 26S proteasome. (**D**) Scheme explaining the counteracting effects of Ubx5 and Rad23 in (A) and (B). When present together, the cofactors achieve a balance between the proteasome and Cdc48 ATPase, which allows efficient protein degradation.

We then asked whether protein degradation would be restored when Ubx5 and Rad23 are both present. Indeed, the inhibition of Ub(n)-FOLD degradation by Rad23 was progressively alleviated by Ubx5 (**Figure 3B**, lanes 6-9), and *vice versa*, inhibition by Ubx5 was reversed by Rad23 (lanes 14-17). Similar results were obtained with Ub(n)-FOLD carrying a shorter polyubiquitin chain (∼5-12 ubiquitin molecules) (**figure S3H**). The degradation of Ub(n)-TAIL was also inhibited by an excess of Ubx5 (**Figure 3A**, lanes 2-5) and restored when Rad23 was also present (lanes 6-9). Adding Rad23 without Ubx5 caused a moderate stimulation of degradation (**Figure 3A**, lanes 10-13), as observed before (**Figure 2G**).

These data indicate that Ubx5 and Rad23 are substrate recruitment factors for Cdc48-UN and the 26S proteasome, respectively, which compete with one another for both folded and unfolded polyubiquitinated substrates (**Figure 3D**). Together, the cofactors achieve a balance between the two molecular machines over a wide concentration range (**Figures 3A** and **3B**), likely by promoting bidirectional substrate shuttling.

### Rad23 and Ubx5 mediate substrate shuttling between the proteasome and Cdc48 complex

To directly determine whether Ubx5 and Rad23 mediate bidirectional substrate shuttling, we first tested whether substrate can be released from the Cdc48-UN complex and transferred to the 26S proteasome (**Figure 4A**). The dissociation of Ub(n)-FOLD from the Cdc48 ATPase complex was determined in the presence of ADP**•** BeF_x_, i.e., under conditions in which substrate cannot be degraded because the ATPases of the 19S regulatory particle are inactive. Polyubiquitinated substrate pre-bound to the Cdc48-UN complex in the presence of Ubx5 could be released by the subsequent addition of 26S proteasomes and Rad23 (**Figure 4A**). Thus, as proposed previously (Finley, 2009), substrates can be transferred from the Cdc48 ATPase to the 26S proteasome.

**Figure 4.**
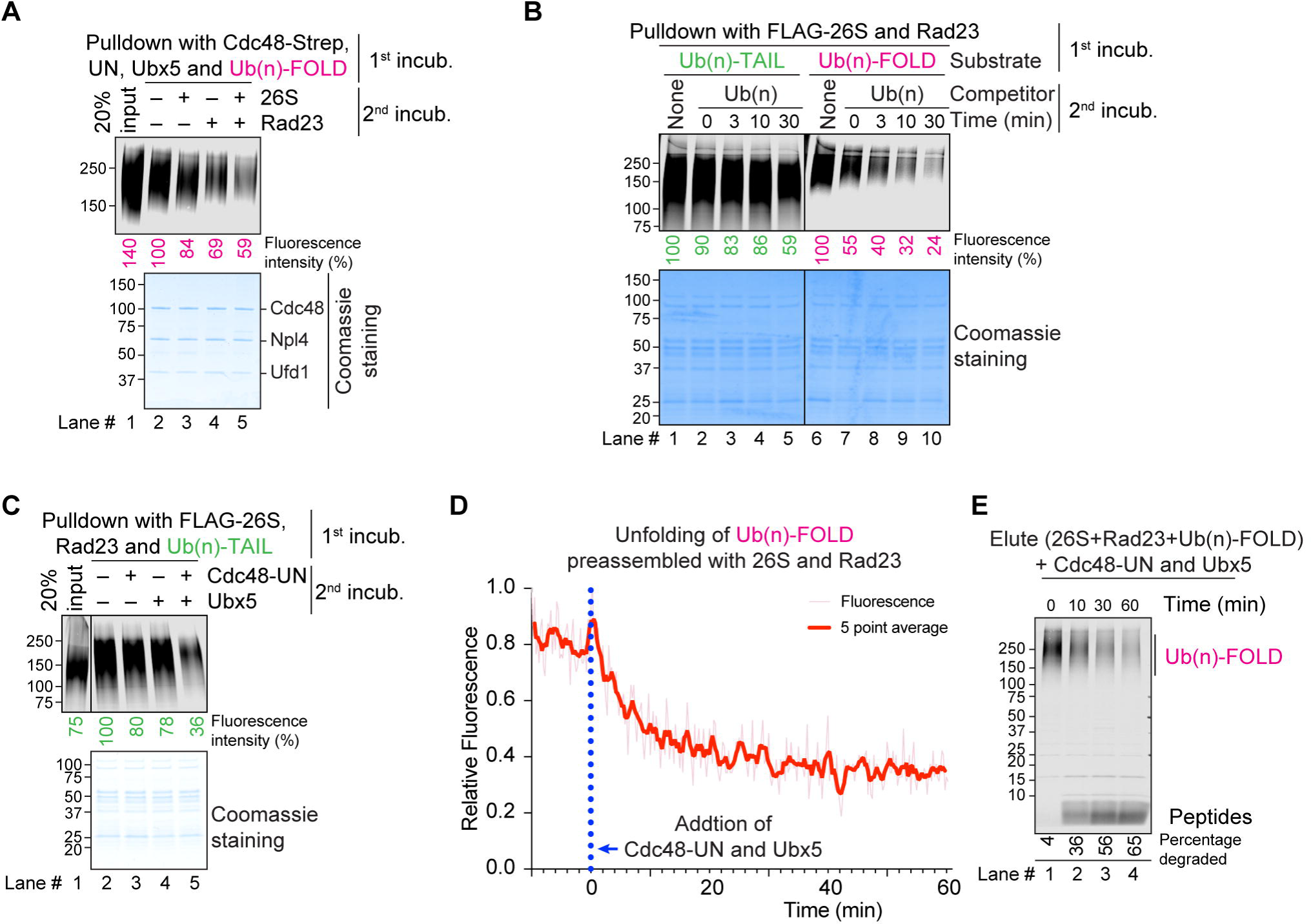
Rad23 and Ubx5 shuttle substrates between the proteasome and Cdc48 ATPase. (**A**) Fluorescently labeled Ub(n)-FOLD was prebound to beads containing streptavidin-tagged Cdc48-UN complex in the presence of ADP• BeF_x_ to prevent ATP hydrolysis-dependent proteolysis (1^st^ incub.). After washing, the beads were incubated with 26S proteasomes and Rad23, as indicated (2nd incub.), and the bound material analyzed by SDS-PAGE, followed by fluorescence scanning and Coomassie-blue staining. Bound substrate was quantified relative to the material bound in the presence of buffer (lane 2 set to 100%). 20% of the input material is shown for comparison (lane 1). (**B**) FLAG-tagged 26S proteasomes were incubated with Rad23 and fluorescently labeled Ub(n)-TAIL or Ub(n)-FOLD in the presence of ADP• BeF_x_ (1^st^ incub.). Proteasomes were retrieved with beads containing FLAG antibodies, washed, and incubated with an excess of free ubiquitin chains (Ub(n)) for different time periods (2^nd^ incub.). The bead-bound material was analyzed by SDS-PAGE, followed by fluorescence scanning and Coomassie-blue staining. Bound substrate was quantified relative to the material bound in the absence of Ub(n) (lanes 1 and 6 set to 100%). (**C**) As in (B), but with the second incubation in the presence of Cdc48-UN complex or Ubx5 instead of free ubiquitin chains. Bound substrate was quantified as in (A). (**D**) FLAG-tagged 26S proteasomes were incubated with Rad23 and fluorescently labeled Ub(n)-FOLD in the presence of ATP. Proteasomes bound to the FLAG beads were eluted and incubated with Cdc48-UN, Ubx5, and ATP. The loss of fluorescence was followed over time. (**E**) As in (D), but samples were taken at different time points and analyzed by SDS-PAGE and fluorescence scanning.

To test whether substrates can be transferred in the reverse direction, we first determined whether polyubiquitinated proteins can spontaneously dissociate from the 26S proteasome/Rad23 complex. Proteasomes were incubated with fluorescent polyubiquitinated substrate and Rad23 (1^st^ incubation), and then an excess of unlabeled free ubiquitin chains (Ub(n)) was added to prevent the re-binding of dissociated substrate (2^nd^ incubation). These experiments showed that the Ub(n)-TAIL substrate was released from proteasomes much slower than Ub(n)-FOLD (**Figure 4B**; lanes 1-5 versus 6-10), consistent with the tail being inserted into the ATPase ring of the 19S particle and providing additional affinity (Bard *et al*., 2019). However, even Ub(n)-TAIL substrate was released from the 26S proteasome-Rad23 complex when both Cdc48-UN and Ubx5 were added during the second incubation (**Figure 4C**). To further prove bidirectional substrate shuttling, we first purified the complex of 26S proteasome, Rad23, and Ub(n)-FOLD by pulling on the proteasome, and then added Cdc48-UN, Ubx5, and ATP. The data show that both substrate unfolding (**Figure 4D**) and degradation (**Figure 4E**) occurred. Unfolding means that Ub(n)-FOLD was transferred from the 26S proteasome to the ATPase complex, and degradation means that, after unfolding, the substrate returned to the proteasome. Together, these results confirm bidirectional shuttling of polyubiquitinated substrates between the 26S proteasome and the Cdc48 ATPase complex.

Next we investigated which domains of Rad23 and Ubx5 are required for substrate shuttling (the domain organizations are shown in **figures S4A**). Rad23 has two UBA domains, one of which is sufficient for interaction with polyubiquitinated substrate (**figure S4A,** lane 5 versus 3). However, both UBA domains are required for inhibition of Ub(n)-FOLD degradation in the absence of Ubx5 (**figure S4B**, lanes 5 and 6 versus 3; **figure S4C**, lane 4 versus 3) and for stimulation of degradation in the presence of Ubx5 (**Figure S4B**, lanes 10 and 11 versus 8; **figure S4C**, lane 7 versus 6). As expected, the UBL domain is also required (**figures S4A; S4B)**. A similar analysis with Ubx5 confirmed that the UBA domain interacts with ubiquitin and the UBX domain with Cdc48 (**figures S4A; S5A**), and showed that both the UBA and UBX are necessary for the inhibition of Ub(n)-FOLD degradation in the absence of Rad23 (**figure S5B,** lanes 4 and 5 versus 3) and the restoration of degradation in the presence of Rad23 (lanes 10 and 11 versus 9). On the other hand, the deletion of the UIM motif or of the UAS domain had no effect (**figure S5B; S5C**). These results show that Rad23 uses its UBA and UBL domains to recruit polyubiquitinated substrates to the 26S proteasome, and Ubx5 uses its UBA and UBX domains to recruit substrates to the Cdc48-UN complex. It should be noted that Rad23 does not interact with Ubx5 with or without added substrate (**figure S5D**, lanes 2 and 3) or the Cdc48-UN complex (**figure S5E**, lanes 2 and 3). Conversely, Ubx5 does not bind to the 26S proteasome/Rad23 complex (**figure S5F**, lanes 2 and 3), and no interaction between the 26S proteasome/Rad23 and Cdc48-UN/Ubx5 complexes was detectable in pull-down experiments, even in the presence of polyubiquitinated substrate (**figure S5G**, lanes 2 and 3). Thus, the two molecular machines exchange substrates, but function independently of each other.

### A role for the Cdc48-cofactor Shp1 in protein degradation

Next we compared Rad23 with the other two UBA-UBL proteins (Dsk2 and Ddi1), and Ubx5 with the other two UBA-UBX proteins (Shp1 and Ubx2). We tested these UBA-UBX and UBA-UBL proteins in all possible combinations in Cdc48-dependent protein degradation by the 26S proteasome, using the model substrates Ub(n)-FOLD and Ub(n)-TAIL (**Figures 5A,B**). The Cdc48-cofactor Shp1 gave the most interesting results.

**Figure 5.**
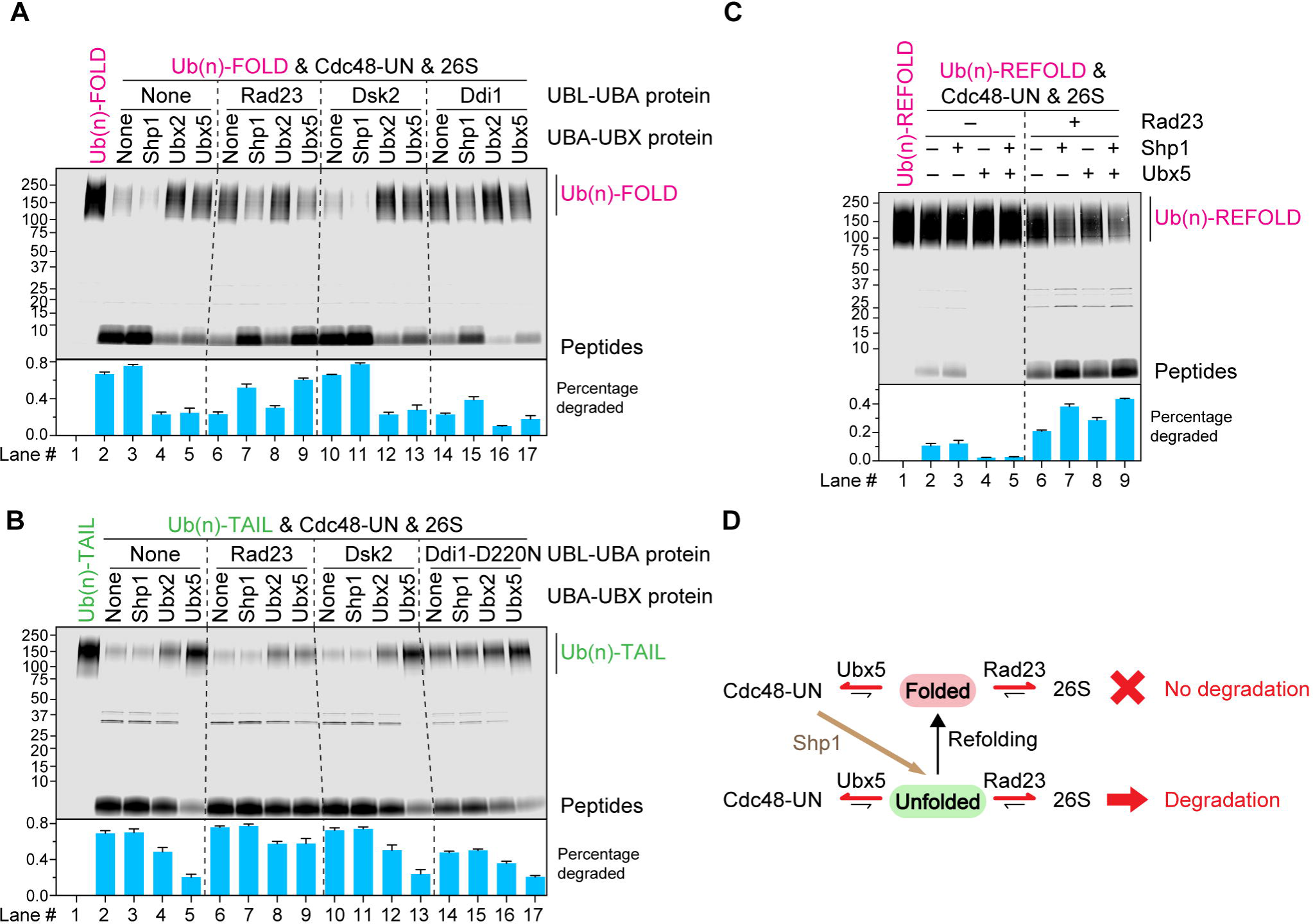
The effect of cofactors on the degradation of different model substrates. (**A**) Fluorescently labeled, photo-converted Ub(n)-FOLD was incubated with 26S proteasomes, Cdc48-UN, the proteasome cofactors Rad23, Dsk2, or Ddi1 (UBA-UBL proteins), and the Cdc48 cofactors Shp1, Ubx2, or Ubx5 (UBA-UBX proteins), as indicated. After 60 min, the samples were analyzed by SDS-PAGE and fluorescence scanning. The percentage of substrate degraded was quantified by determining the fluorescence in peptides (after background subtraction) and comparing it to the total fluorescence in each lane of the SDS gel. The experiment was performed in triplicates and the means and standard deviations are shown. (**B**) As in (A), but with Ub(n)-TAIL. (**C**) Ub(n)-REFOLD, a substrate that spontaneously refolds after Cdc48-mediated unfolding (**figure S7E**), was used in degradation experiments with the indicated components. Quantification was done as in (A). (**D**) Scheme explaining the effect of cofactors on the degradation of Ub(n)-REFOLD in (E). Substrate refolding (arrow) competes with degradation. Ubx5 recruits folded substrate to Cdc48-UN, Shp1 stimulates substrate unfolding, and Rad23 recruits unfolded substrate to the proteasome.

When tested with Ub(n)-FOLD, Shp1 was able to relieve the inhibition caused by Rad23 (**Figure 5A**, lane 7 versus 6), similarly to Ubx5 (lane 9 versus 6). However, Shp1 did not inhibit the degradation of the folded substrate in the absence of Rad23 and was in fact slightly stimulatory (**Figure 5A**, lane 3). Together with the pull-down experiments (**figure S3F**), these results indicate that Shp1 does not stimulate degradation by promoting substrate binding to the Cdc48 complex. Rather, it seems to promote the unfolding of substrate molecules that already are bound to the Cdc48 complex. Consistent with this assumption, Shp1, in contrast to Ubx5, stimulated the unfolding of Ub(n)-FOLD (**figure S3G**) and had little effect on the degradation of Ub(n)-TAIL (**Figure 5B**, lane 3 versus 2), a substrate that does not need prior unfolding.

A domain analysis showed that Shp1 without its UBA domain lost the interaction with the ubiquitin chain of the substrate **(figure S6A,** lane 3 versus 2), but still relieved the inhibition of Ub(n)-FOLD degradation by Rad23 (**figure S6B,** lanes 8 and 9 versus 7). These data thus confirm that Shp1 does not function by promoting substrate recruitment to the Cdc48-UN complex. Nevertheless, an interaction of Shp1 with Cdc48 is important for counteracting Rad23, as the deletion of the UBX domain abolished the stimulatory effect of Shp1 on substrate degradation (**figure S6B,** lanes 11 versus 8). The SEP domain is also required (lane 10 versus 8).

Interestingly, Shp1 did not counteract Rad23 without the UN cofactor (**figure S6C,** lane 9 versus 8), suggesting that Shp1 and UN must both interact with Cdc48 to stimulate the degradation of Ub(n)-FOLD. Whereas Shp1 bound to Cdc48 in the absence of UN (**figure S6D**, lane 5), only small amounts associated with Cdc48-UN even in the presence of substrate (**figure S5A,** lane 3; Coomassie-stained gel and anti-SBP blot), indicating that the two Cdc48 cofactors do not bind at the same time, as reported for the mammalian homologs (Bruderer *et al*., 2004). Nevertheless, Shp1 moderately stimulated the ATPase activity of Cdc48-UN (**figure S6E**), suggesting that the cofactors can in fact cooperate. Taken together, our results suggest that Shp1 functions by activating the Cdc48-UN complex, enhancing the generation of unfolded substrate molecules for subsequent degradation by the 26S proteasome.

### The other proteasome and Cdc48 cofactors do not promote protein degradation

The remaining UBA-UBL proteins (Ddi1 and Dsk2) and the UBA-UBX protein Ubx2 did not promote Cdc48-dependent protein degradation by the 26S proteasome. The UBA-UBL protein Ddi1 was inhibitory in all our assays (**Figures 5A,B**, lane 14 versus 2; **figure S3C**), likely because it strongly interacts with the polyubiquitin chain (Yip *et al*., 2020) and therefore prevents the substrate from associating with the proteasome and Cdc48 complex.

The UBA-UBX protein Ubx2 was also mostly inhibitory (**Figures 5A,B**, lane 4 versus 2; **figures S3G**; **S6E**). Because Ubx2 stimulates substrate recruitment to Cdc48-UN (**figure S3F**), it seems to sequester substrate on the ATPase complex and keep it away from the proteasome, explaining why Ubx2 inhibits the degradation of Ub(n)-FOLD more than that of Ub(n)-TAIL (**Figures 5A; 5B**).

Dsk2 was inactive in all our assays (**Figures 5A,B**, lanes 10-13 versus 2-5; **Figure 2B**, lane 4 versus 2; **figure S3C**). Thus, Dsk2 behaves differently from Rad23, although both proteins have overlapping functions *in vivo* (Biggins et al., 1996; Kim et al., 2004; Medicherla *et al*., 2004). A chimera containing the UBL domain of Dsk2 and the two UBA domains of Rad23 was only partially active (**figure S7A**, lane 5 versus 3 and lane 9 versus 7), and replacement of the UBL domain of Dsk2 with that of Rad23 rendered the protein inactive (lanes 4 and 8). Thus, Dsk2 seems to be inactive because it lacks a second UBA domain and has a suboptimal UBL domain. We wondered if the single UBA domain would be sufficient if the substrate carried two ubiquitin chains. Such a substrate, designated Ub(n)2-FOLD, was generated by a protocol similar to one previously described (Blythe et al., 2017) (**figure S7B**) and contained a cleavage site for the TEV protease, which allowed us to ascertain that the substrate indeed contained two ubiquitin chains (**figure S7C**). However, Dsk2 was still inactive (**figure S7D**, lanes 4-6 versus 3, and 9-11 versus 8).

### Cdc48-dependent degradation of a rapidly refolding substrate

Given that both model substrates Ub(n)-TAIL and Ub(n)-FOLD could be degraded in the absence of Rad23, Ubx5, and Shp1 (**Figures 3A; 3B**), we wondered what advantage these cofactors might provide. One possibility is that they are required for substrates that escape efficient degradation because they spontaneously refold after processing by Cdc48. We therefore generated the substrate Ub(n)-REFOLD by omitting photocleavage; this substrate thus differs from Ub(n)-FOLD by having an intact Dendra moiety. As observed with a similar substrate (Ji *et al*., 2022), Ub(n)-REFOLD rapidly refolded after its processing by the Cdc48 ATPase, resulting in Dendra fluorescence remaining almost constant (**figure S7E**). Ub(n)-REFOLD was not efficiently degraded when only 26S proteasomes and Cdc48-UN were added (**Figure 5C**, lane 2), in contrast to the results with Ub(n)-TAIL and Ub(n)-FOLD (**Figures 1B; 1D**). Shp1 had a slight stimulatory effect when added alone (**Figure 5C**, lane 3), whereas Ubx5 was inhibitory (lanes 4 and 5). However, when Rad23 was added together with either Shp1 or Ubx5, the degradation of Ub(n)-REFOLD was stimulated (lanes 7 and 8). Degradation was most efficient when all cofactors were present (lane 9). These results support a model in which Ubx5 recruits folded protein molecules to the Cdc48 ATPase for their unfolding, Shp1 stimulates the Cdc48 ATPase to generate more unfolded protein molecules, and Rad23 recruits unfolded protein molecules back to the proteasome for their degradation, thereby counteracting the rapid refolding of the substrate (**Figure 5D**).

We also tested the two substrates with suboptimal flexible segments (Ub(n)-NTAIL and Ub(n)-CTAIL), which were inefficiently degraded by the 26S proteasome alone (**figures S2D; S2F**), in the presence of Cdc48-UN and the cofactors. Ub(n)-NTAIL behaved similarly to Ub(n)-REFOLD (**figure S7F**), likely because the Dendra protein fused to the unstructured segment had to be transiently unfolded by Cdc48-UN complex to provide additional initiation segments for the proteasome. The Ub(n)-CTAIL substrate was degraded when Rad23 was added (**figure S7G,** lane 3), indicating that Rad23 promotes the proper insertion of the flexible segment into the 26S proteasome. Rad23 counteracted the inhibition exerted by Cdc48-UN alone (**figure S7G,** lane 5 versus 4), but addition of Shp1 and Ubx5 re-established the inhibition (lane 6). Thus, this substrate might require additional proteasome-recruitment factors for efficient degradation.

### Bidirectional shuttling of polyubiquitinated substrates *in vivo*

Finally, we tested whether bidirectional substrate shuttling between the 26S proteasome and Cdc48 ATPase complex occurs *in vivo.* We used a temperature-sensitive *S. cerevisiae* mutant (*cdc48-3*), in which folded proteins are inefficiently degraded because their unfolding by the Cdc48 ATPase is impaired. The growth defect of *cdc48-3* cells at elevated temperatures was partially relieved when Ubx5 was overexpressed (**Figure 6A**). Shp1 overexpression had no effect. Thus, Ubx5 seems to increase the efficiency of substrate recruitment to the Cdc48 mutant. As reported (Tsuchiya *et al*., 2017; Verma et al., 2011), polyubiquitinated proteins accumulated on FLAG-tagged 26S proteasomes in *cdc48-3* cells, but not in wild-type cells (**Figure 6B**, lanes 4-6 versus 1-3) or in *cdc48-3* cells lacking Rad23 and Dsk2 (lanes 13-15). When Ubx5 was overexpressed in *cdc48-3* cells, the amount of proteasome-associated polyubiquitinated proteins decreased more than two-fold (lanes 10-12). Polyubiquitinated proteins also accumulated on the Cdc48-UN complex in cdc48-3 cells compared to wild-type cells, as determined by pull-down experiments with HA-tagged Npl4 (**Figure 6C**, lanes 4-6 versus 1-3). Less accumulation was seen in the absence of Ubx5 (*ubx5Δ* cells) (lanes 7-9) and more when Ubx5 was overexpressed (lanes 10-12). These data are consistent with Ubx5 moving polyubiquitinated proteins from the 26S proteasome/Rad23 complex to the Cdc48 ATPase complex.

**Figure 6.**
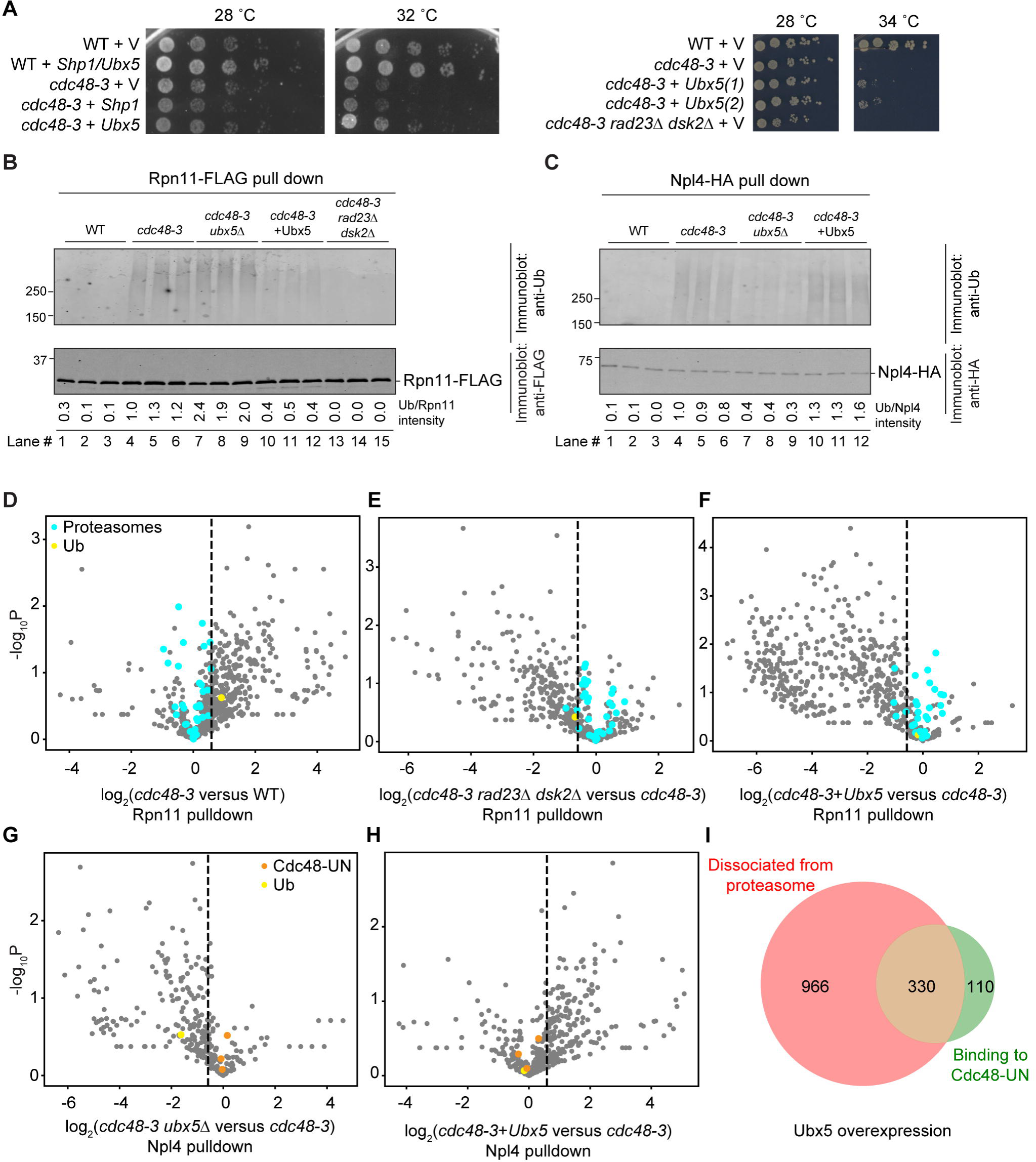
Rad23 and Ubx5 compete for polyubiquitinated substrates *in vivo*. (**A**) Wild-type (WT) or cdc48-3 cells were transformed with a vector (V) or a 2μ plasmid expressing SBP-tagged Ubx5 or Shp1 under the endogenous promoters. The cells were plated after serial dilution and incubated at 28°C, 32°C, or 34 °C. (**B**) The indicated strains expressed the FLAG-tagged proteasomal subunit Rpn11 and HA-tagged Npl4. The proteasomes were immunoprecipitated from cell extracts with anti-FLAG beads and the bound material analyzed by SDS-PAGE, followed by immunoblotting with ubiquitin and FLAG antibodies. The amounts of bound ubiquitinated proteins were quantified and normalized relative to the intensity of the Rpn11-FLAG band. (**C**) As in (B), but Cdc48-UN was immunoprecipitated with anti-HA beads. (**D**) Mass spectrometry analysis comparing proteasome-associated proteins from cdc48-3 cells with those from WT cells. Each point in the volcano plot represents a protein for which the ratio of its abundance in the mutant and in WT cells is given, as well as a measure of statistical significance (P-value), derived from a multiple non-parametric t-test based on three biological replicates. Proteasome subunits are labeled in cyan and ubiquitin in yellow. Proteins to the right of the dashed line are upregulated by a factor of >1.5. (**E**) As in (D), but comparing proteasome-associated proteins from cdc48-3 cells with those in cdc48-3 cells lacking Rad23 and Dsk2. (**F**) As in (D), but comparing proteasome-associated proteins from cdc48-3 cells with those in cdc48-3 cells overexpressing Ubx5. (**G**) As in (D), but comparing Cdc48-associated proteins from cdc48-3 cells with those from WT cells. (**H**) As in (D), but comparing Cdc48-associated proteins from cdc48-3 cells with those in cdc48-3 cells overexpressing Ubx5. (**I**) Overexpression of Ubx5 in *cdc48-3* cells moves a large number of proteins from the proteasome to the Cdc48 complex (overlapping region of the diagram; see also Table S2).

To identify specific substrates that shuttle between the proteasome and Cdc48 ATPase, we performed proteomics experiments. We first used mass spectrometry to identify proteins that accumulate on 26S proteasomes in *cdc48-3*, but not wild-type cells (**Figure 6D; Table S1**). These are likely folded proteins that require efficient Cdc48-mediated unfolding for their proteasomal degradation. As expected, most of these proteins no longer associated with the proteasome when Rad23 and Dsk2 were absent (**Figure 6E**). Ubx5 overexpression also caused the dissociation of a large number of proteins from the proteasome (**Figure 6F**). Consistent with the immunoblotting experiments (**Figures 6C**), many proteins did not bind to the Cdc48-UN complex in the absence of Ubx5 (**Figures 6G**) and accumulated on the complex when Ubx5 was overexpressed (**Figures 6H**). Importantly, Ubx5 overexpression caused many of the proteins released from the 26S proteasome to associate with the Cdc48-UN complex (**Figures 6I; Table S2**). This population may be an underestimate because the pull-down of the ATPase complex was less efficient than that of the proteasome. Nevertheless, approximately 75% of the proteins accumulating on the Cdc48-UN complex originate from the proteasome. The list of shuttling substrates contains proteins of different cellular localizations (**Table S2**). Taken together, these data indicate that a broad range of proteins shuttle between the 26S proteasome and the Cdc48 ATPase complex in a cofactor-dependent manner.

## DISCUSSION

Here we have reconstituted Cdc48-dependent degradation of polyubiquitinated model substrates by the proteasome, using purified components from *S. cerevisiae*. We show that a minimal system that allows the efficient degradation of proteins with different folding characteristics consists of the 26S proteasome, the Cdc48-UN ATPase complex, the proteasome cofactor Rad23, and the Cdc48 cofactors Ubx5 and Shp1. Based on our results, we propose a novel model for how the 26S proteasome and Cdc48 ATPase complex cooperate in protein degradation (**Figure 7**). Rad23 and Ubx5 serve as recruitment factors for the 26S proteasome and the Cdc48-UN complex, respectively, allowing the two molecular machines to compete for polyubiquitinated substrates before and after their unfolding. Shp1 participates by stimulating protein unfolding by the Cdc48-UN complex. Our *in vivo* experiments show that a large number of cellular proteins require bidirectional substrate shuttling between the 26S proteasome and Cdc48 ATPase for their degradation.

**Figure 7.**
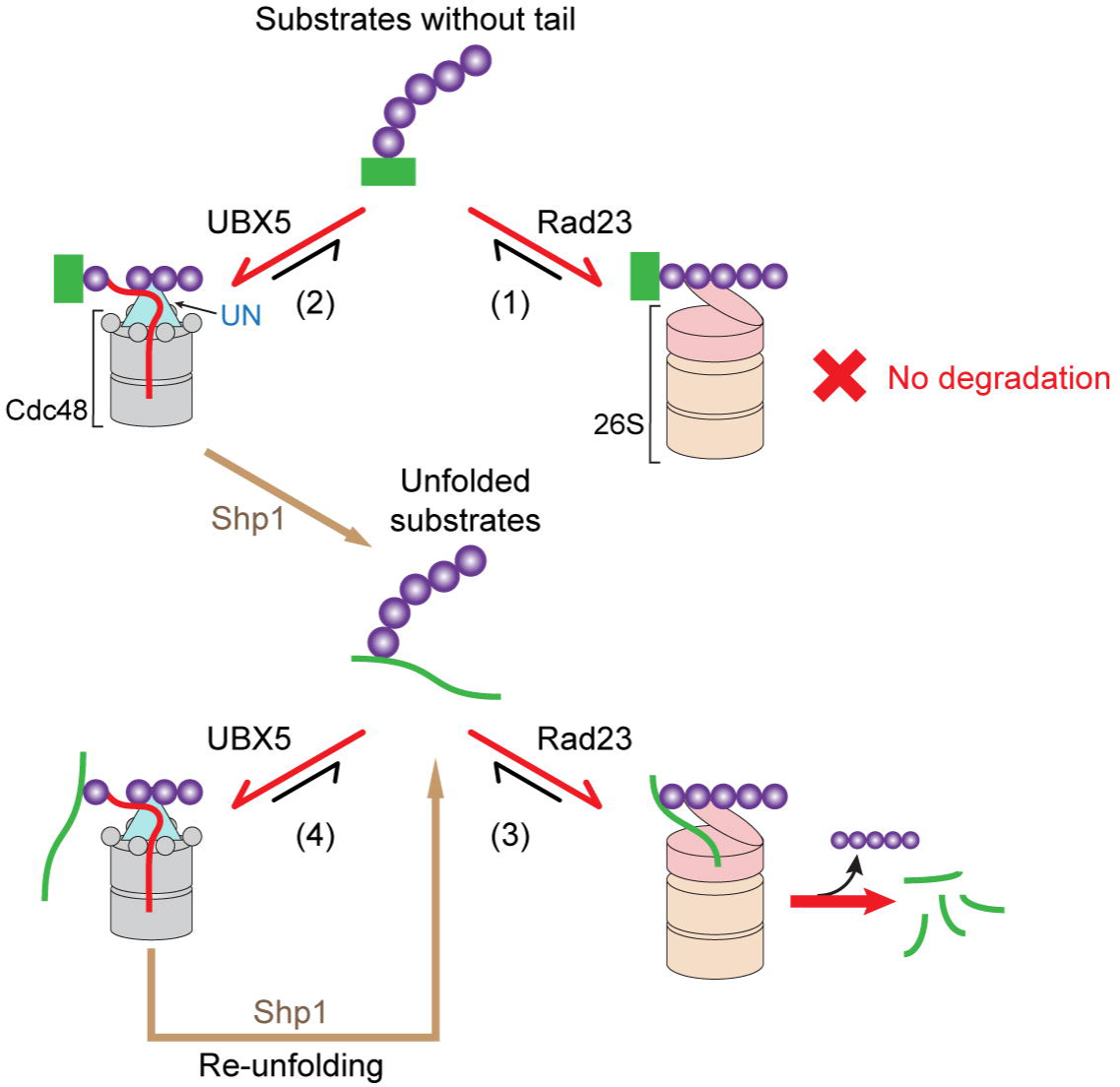
Model for the interplay of the 26S proteasome and Cdc48 ATPase complex in protein degradation. For details, see text.

The 26S proteasome seems to be the major component that discriminates between folded and unfolded substrates, as it can only degrade proteins with an unstructured segment; the other components, i.e. the Cdc48-UN complex, Rad23, Ubx5, and Shp1, primarily interact with the ubiquitin chain. When a well-folded protein is recruited by Rad23 to the 26S proteasome, it would not be degraded, as it lacks a flexible segment that could initiate translocation into the proteolytic chamber (**Figure 7**; reaction 1). The substrate would then dissociate, bind through Ubx5 to the Cdc48-UN complex (reaction 2), and be unfolded by the Cdc48 ATPase in a process that is stimulated by Shp1. Alternatively, the folded substrate could bind first to the ATPase complex and be unfolded, without initially encountering the proteasome. After unfolding, the proteasome and Cdc48 ATPase complex would again compete with one another for the polyubiquitinated substrate (**Figure 7**, reactions 3 and 4). Recruitment to the proteasome would cause its degradation (reaction 3), as it now contains a flexible segment. The substrate may instead rebind to the Cdc48 complex resulting in another round of translocation through the central pore of Cdc48 (reaction 4). Unfolding by Cdc48 could be repeated, but because the proteasome competes with the ATPase during each cycle, ultimately all substrate molecules would be degraded. Substrates that already contain a flexible tail could be degraded directly by the 26S proteasome. Because tail insertion retards substrate dissociation from the proteasome, transfer to the Cdc48 ATPase would be disfavored, so that many proteins would be degraded without encountering the Cdc48 ATPase. If proteins with a flexible segment bind first to the Cdc48 complex, their degradation would be delayed, particularly if the segment is suboptimal for recruitment to the proteasome and the remainder of the substrate can rapidly refold.

The conclusions from our *in vitro* reconstitutions are supported by *in vivo* experiments. Our results agree with reports in the literature that Rad23, Ubx5, and Shp1 are involved in the degradation of various substrates in *S. cerevisiae* (Lambertson *et al*., 1999; Rao and Sastry, 2002; Schuberth et al., 2004; Verma *et al*., 2011), and that Rad23 and Shp1 participate in ERAD (Medicherla *et al*., 2004; Tran et al., 2011). A role for Cdc48 acting downstream of the proteasome is also consistent with previous data showing that Cdc48 and the proteasome can compete for polyubiquitinated substrates *in vivo* (Verma *et al*., 2011) and that Cdc48 can mobilize a fragment of a transcription factor generated by partial proteasomal cleavage (Rape et al., 2001).

Our data show that the combination of 26S proteasome and Cdc48-UN complex is sufficient to degrade a substrate with flexible segments and a folded substrate that is irreversibly unfolded by the Cdc48 ATPase. Such a rudimentary system does not allow the efficient degradation of substrates that spontaneously refold after Cdc48-mediated unfolding, an observation that may explain the low efficiency of a previous reconstitution system (Olszewski *et al*., 2019). Of course, a system consisting of only 26S proteasomes and Cdc48-UN does not normally exist in cells, but its ability to degrade many proteins may explain why the UBA-UBL and UBA-UBX proteins are dispensable for the viability of yeast cells under normal growth conditions. Surprisingly, we found a substrate with a flexible segment, (Ub(n)-CTAIL), that is not efficiently degraded in the presence of Cdc48 complex and all cofactors. The poor degradation efficiency might be due to a combination of a suboptimal distance between the ubiquitin chain and the unstructured segment and the ability of the remainder of the substrate to rapidly refold. It remains unclear whether such substrates are poorly degraded *in vivo* or use additional proteasome-recruitment factors.

Previous models considered proteasome-recruitment factors, such as Rad23, to mediate substrate shuttling from the Cdc48 ATPase to the proteasome. Our results now show that Rad23 alone would favor the direct recruitment of folded substrates to the proteasome, where they could neither be degraded nor be transferred to Cdc48-UN. Rad23 therefore needs to cooperate with the Cdc48-cofactors Ubx5 or Shp1, whose role in protein degradation had been unclear. Our results show that Cdc48 cofactors function in a hierarchical manner: the UN cofactor is the basic one, while Ubx5 and Shp1 associate subsequently. Ubx5 acts as a substrate recruitment factor for the Cdc48-UN complex in a similar way as Rad23 for the proteasome. Both bind to their respective partners through ubiquitin-like domains: Rad23 associates with the 26S proteasome through its UBL domain and Ubx5 with Cdc48 through its UBX domain. In both cases, the binding seems to be synergistic with polyubiquitinated substrate, which might be explained by avidity effects or by the relief of auto-inhibition caused by an intramolecular interaction between the UBA and ubiquitin-like domains (Kochenova et al., 2022; Ryu et al., 2003). Shp1 is not a recruitment factor, but seems to accelerate the rate of substrate unfolding, consistent with the mammalian homologs stimulating the ATPase activity of p97 (Zhang et al., 2015). Surprisingly, Shp1’s function is dependent on the UN complex, although only a small amount of Shp1 associates with Cdc48-UN. We speculate that UN and Shp1 bind to Cdc48 at different stages during substrate processing, UN during initiation and Shp1 during subsequent polypeptide translocation through the pore. *In vivo*, Shp1 and Ubx5 probably function together, as they can act synergistically in our *in vitro* system. However, Ubx5 might deal with shorter ubiquitin chains than Shp1 (**figure S3B**, lane 8 versus 6), consistent with data on UbxN7, the mammalian homolog of Ubx5 (Fujisawa et al., 2022).

The third UBA-UBX protein Ubx2 does not seem to allow substrate shuttling from the Cdc48 complex to the proteasome. Ubx2 also differs from Ubx5 and Shp1 by being bound to membranes (Schuberth and Buchberger, 2005). Therefore, Ubx2 may not promote the degradation of cytosolic proteins, but rather recruit the Cdc48-UN complex to membranes, for example, during ERAD.

We were surprised that the UBA-UBL protein Dsk2 was inactive in our *in vitro* system; *in vivo,* Dsk2 and Rad23 have redundant functions (Biggins et al., 1996; Kim et al., 2004; Medicherla *et al*., 2004). Dsk2 differs from Rad23 by lacking a second UBA domain and possessing a suboptimal UBL domain, which explains why it was inactive in our system. Our data indicate that Dsk2 cannot recruit substrates to the proteasome exclusively through the attached ubiquitin chain; even two ubiquitin chains were insufficient. *In vivo*, Dsk2 likely uses its UBA domain to bind to ubiquitin and its middle domain to bind to polypeptide segments of the substrate, as demonstrated for the mammalian homolog (Itakura et al., 2016).

The UBA-UBL protein Ddi1 also did not bind to 26S proteasomes, consistent with its weak binding to Rpn1 (Gomez et al., 2011; Rosenzweig et al., 2012). It remains unclear whether Ddi1 functions as a shuttling factor, like its mammalian homolog Ddi2 (Collins et al., 2022), or acts as an independent protease that cleaves substrates with very long ubiquitin chains (Yip *et al*., 2020).

Our results likely apply to higher eukaryotes, as a general role for the mammalian Rad23 homologs HR23A and HR23B in proteasomal protein degradation has been well established (Yokoi and Hanaoka, 2017), and the Ubx5 homolog Ubxn7 is involved in the degradation of several proteins, including HIF1α, NRF2, and components of the DNA replication machinery (Alexandru et al., 2008; Di Gregorio et al., 2021; Tarcan et al., 2022). The role of the mammalian Shp1 homologs p47 and p37 is less clear, as they have been implicated mostly in the ubiquitin-independent disassembly of protein complexes by the p97 ATPase (Kracht et al., 2020; van den Boom et al., 2021; Weith et al., 2018). However, p47 has been reported to have a role in ERAD (Park et al., 2017), suggesting that it may participate in ubiquitin-dependent proteasomal degradation, similar to Shp1 in *S. cerevisiae* (Schuberth *et al*., 2004).

### Limitations of the study

Future work needs to clarify the mechanism by which Shp1 stimulates substrate processing by the Cdc48-UN complex. Specifically, the hypothesis should be tested that Shp1 acts at a step after substrate recruitment by UN. The role of DUBs in the shuttling of substrates between the 26S proteasome and Cdc48-UN also needs to be explored. Because Cdc48 and the 26S proteasome differ slightly in their requirement of a minimum ubiquitin chain length (five versus four), a DUB could potentially change the balance between the two molecular machines. Although all tested components have homologs in mammals, the Cdc48 homolog p97 has additional cofactors that could have a role in substrate shuttling between the ATPase and the proteasome.

## ACKNOWLEDGEMENTS

We thank Ellen A. Goodall, Michael L. Skowyra, Daniel Finley, Alfred L. Goldberg, Galen A. Collins, Donghoon Lee, and Nicholas O. Bodnar for critical reading of the manuscript. We are particularly grateful to Michael L. Skowyra for helpful comments. This work was supported by a NIGMS grant (R01 GM052586) to T.A.R. Z.J. was a Howard Hughes Medical Institute Fellow of

the Damon Runyon Cancer Research Foundation, DRG-2315-18. T.A.R. is a Howard Hughes Medical Institute Investigator.

## AUTHOR CONTRIBUTIONS

H.L. performed all experiments, J.A.P. and S.P.G. performed the mass spectrometry analysis, T.A.R. supervised the project. H.L., Z.J., and T.A.R. wrote the manuscript.

## STAR * METHODS

### RESOURCE AVAILABILITY

#### Lead Contact

Further information and requests for resources and reagents should be directed to and will be fulfilled by the Lead Contact, Tom Rapoport.

#### Materials Availability

All unique/stable reagents generated in this study are available from the Lead Contact with a completed Materials Transfer Agreement.

#### Data and Code Availability

- This paper does not report original code.
- Any additional information required to reanalyze the data reported in this paper is available from the lead contact upon request.

### EXPERIMENTAL MODEL AND SUBJECT DETAILS

#### Yeast strains and cultures

Plasmids encoding *S. cerevisiae* Uba1, Ubr1, and Ubx2 were transformed into the INVSc1 yeast strain (Thermo Fisher). Yeast cells were grown in synthetic dropout (SD) medium for 24 h, and then switched to medium containing 2% galactose to induce protein expression. The cells were harvested after 24 h of galactose induction.

Yeast strain YYS40 (genotype *MAT*a *ade2-1 his3-11,15 leu2-3,112 trp1-1 ura3-1 can1 Rpn11::Rpn11-*3×Flag) used for proteasome purification was a gift from Andreas Martin (UC Berkeley). The cells were grown at 30°C in YPD for 2 days until saturation. Yeast strain sDL135 (genotype *MAT*a *lys2-801 leu2-3, 2-112 ura3-52 his3-Δ 200 trp1-1 pre1::PRE1-TEVProA (HIS3)*) used for 20S proteasome purification was a gift from Daniel Finley (Harvard Medical School). The cells were grown in YPD for 24 h at 25°C and another 24 h at 20°C. Yeast strains 3592-4-4 (*Mat*a *pdr5::KanMX RPN11-3×FLAG::HIS3*), 3592-5-2 (*Mat*a *pdr5::KanMX cdc48-3 RPN11-3×FLAG::HIS3*), and 3967-2-4 (*Mat*a *pdr5:Kan cdc48-3 dsk2::TRP1 rad23::Sphis5 +RPN11-3×FLAG::HIS3*) were gifts from Yanchang Wang (Florida State University). Yeast strains *Mat*a *pdr5::KanMX RPN11-3×FLAG::HIS3 NPL4-3xHA::LEU2*, *pdr5::KanMX cdc48-3 RPN11-3×FLAG::HIS3 NPL4-3xHA::LEU2*, *pdr5::KanMX cdc48-3 RPN11-3×FLAG::HIS3 NPL4-3xHA::LEU2 ubx5::URA3*, *Mat*a *pdr5:Kan cdc48-3 dsk2::TRP1 rad23::Sphis5 +RPN11-3×FLAG::HIS3 NPL4-3xHA::LEU2* were generated by homologous recombination. Plasmids encoding *Saccharomyces cerevisiae* Shp1 or Ubx5 were transformed into the 3592-5-2 or *pdr5::KanMX cdc48-3 RPN11-3×FLAG::HIS3 NPL4-3xHA::LEU2* strain. The cells were grown in SD medium at 25°C to OD_600_ of 0.6 and shifted to 34°C for another 1 h.

#### Bacterial cultures

Plasmids were transformed into *Escherichia coli* BL21 CodonPlus (DE3) RIPL cells (Agilent), unless stated otherwise. The cells were grown in Terrific Broth to an OD_600_ of 0.8. Protein expression was induced by addition of 0.1 mM isopropyl b-D-1-thiogalactopyranoside (IPTG), and the incubation was continued at 16°C for 16 h.

### METHOD DETAILS

#### Plasmids

*S. cerevisiae* ubx2 was cloned into the pRS426Gal1 vector with an N-terminal modified His14-tag (HHHHSGHHHTGHHHHSGSHHH) followed by a Tobacco Etch Virus (TEV)-protease cleavage site (ENLYFQG), and a C-terminal HRV 3C protease cleavage site (LEVLFQG) followed by a streptavidin-binding protein (SBP) tag (MDEKTTGWRGGHVVEGLAGELEQLRARLEHHPQGQREP). *S. cerevisiae* Shp1 and Ubx5 were cloned into the pRS425 and pRS426 vectors, respectively, each contains its endogenous promoter and terminator with a C-terminal HRV 3C protease cleavage site followed by a SBP tag.

Wild-type Cdc48 and its variants were cloned into the pET28 vector using NotI and AscI sites, with a His6-tag and a TEV-protease cleavage site at the N-terminus. For Cdc48 pull-down experiments, a HRV 3C protease cleavage site followed by two copies of a sequence encoding the strep tag (WSHPQFEK) was added at the C-terminus of Cdc48. All Ufd1 variants were cloned into the pK27 vector with an N-terminal His14-SUMO (small ubiquitin-like modifier) tag. A sequence encoding the hemagglutinin (HA)-tag (YPYDVPDYA) was inserted between SUMO and Ufd1 where indicated. All Npl4 variants were cloned into the pET21 vector using NdeI and AscI sites, with a C-terminal His6-tag or FLAG-His6 (DYKDDDDKGLEHHHHHH) tag.

Wild-type Rad23, Dsk2, Ddi1, Shp1, Ubx5 and their variants were cloned into the pET28 vector using NotI and AscI sites, with N-terminal His6-tag and a TEV-protease cleavage site, and a C-terminal HRV 3C cleavage site followed by a SBP tag. In some experiments, Rad23 and Ubx5 were used without 3C site and SBP tag.

A bacterial expression plasmid coding for an N-end degron (NeD) sequence with lysine-less super-folder GFP (NeD-sfGFP) has been described (Ji *et al*., 2022). This construct was used to generate Ub(n)-TAIL. The folded substrate (Ub-G76V-Dendra) was constructed by fusing ubiquitin with a C-terminal G76V mutation to Dendra, with a short linker (GSCGSGGS) in between. A His6-tag followed by a TEV-protease cleavage site was added to the N-terminus. The construct was generated by overlapping PCR and then cloned into the pET28 vector using NotI and AscI sites. This construct was used to generate Ub(n)-FOLD, Ub(n)-REFOLD, and Ub(n)-FOLD(s). The constructs for Ub(n)-NTAIL (NTAIL-Ub-G76V-Dendra) and Ub(n)-CTAIL (Ub-G76V-Dendra-CTAIL) were generated from Ub-G76V-Dendra by adding either the NeD sequence at the N-terminus or a SBP tag at the C-terminus. The construct for Ub(n)2-FOLD (Ub-G76V-Ub-G76V-Dendra) was generated by addition of another Ub-G76V at the N-terminus.

The bacterial expression plasmid encoding *S. cerevisiae* Ubc2 has been described in (Bodnar and Rapoport, 2017b). The bacterial expression plasmids encoding gp78^RING^-Ube2g2 and His14-SUMO-hUb, and yeast expression plasmids endocing *S. cerevisiae* Uba1 and Ubr1 have been described in (Ji *et al*., 2022). A plasmid encoding mouse Ube1 was a gift from Jorge Eduardo Azevedo (Addgene plasmid # 32534).

#### Immunoblotting and antibodies

Antibodies used in this study were: anti-SBP-tag (Millipore, clone 20, 1:1000), anti-FLAG antibody (Millipore, 1:1000), human ubiquitin antibody (R&D Systems, 1:2000), anti-HA antibody (Millipore, 1:2000), anti-Cdc48 antibody (MyBioSource, 1:500), donkey anti-mouse IgG DyLight 680 conjugated (Thermo Fisher, 1:5000), donkey anti-rabbit IgG DyLight 680 conjugated (Thermo Fisher, 1:5000), donkey anti-mouse IgG DyLight 800 conjugated (Thermo Fisher, 1:5000), donkey anti-rabbit IgG DyLight 800 conjugated (Thermo Fisher, 1:5000), goat anti-rat IgG DyLight 800 conjugated (Thermo Fisher, 1:5000), and donkey anti-goat IgG DyLight 680 conjugated (Thermo Fisher, 1:5000).

#### Protein purifications

Cdc48 and its variants, untagged Ufd1/Npl4 (UN), HA-Ufd1, Npl4-FLAG, and NeD-sfGFP were expressed and purified as previously described (Ji *et al*., 2022). Rad23, Dsk2, Ddi1, Shp1, Ubx5, and their variants were purified as follows. Bacterial cells were harvested by centrifugation at 5,000 x g for 10 min and resuspended in wash buffer (50 mM Tris-HCl, pH 8, 320 mM NaCl, 5 mM MgCl_2_, 10 mM imidazole, 0.5 mM ATP) supplemented with phenylmethylsulfonyl fluoride (PMSF; 1 mM) and a protease inhibitor cocktail. The cells were lysed by sonication. The lysate was cleared by ultracentrifugation in a Ti-45 rotor (Beckman) at 40,000 rpm for 30 min at 4°C. The supernatant was incubated for 60 min at 4°C with Ni-NTA resin that was pre-equilibrated with wash buffer. The resin was washed three times with 30 column volumes of wash buffer. Protein was eluted with elution buffer (50 mM Tris-HCl, pH 8, 150 mM NaCl, 5 mM MgCl_2_, 400 mM imidazole), and the eluate was concentrated and subjected to size-exclusion chromatography (SEC) on a Superdex 200 Increase column, equilibrated in SEC buffer (50 mM HEPES, pH 7.4, 150 mM NaCl, 5 mM MgCl_2_, and 0.5 mM tris(2-carboxyethyl)phosphine (TCEP)). The purified protein was snap-frozen in SEC buffer.

The constructs for Ub(n)-FOLD (Ub-G76V-Dendra) and Ub(n)2-FOLD (Ub-G76V-Ub-G76V-Dendra) was purified by Ni-NTA resin as described above. TEV protease was added to the eluted proteins and 10 mM imidazole was added. The samples were then incubated with Ni-NTA resin to remove the His tag, and the unbound protein was concentrated and loaded onto a Superdex 200 Increase column equilibrated with SEC buffer. Ub-G76V-Dendra and Ub-G76V-Ub-G76V-Dendra used for Ub(n)-FOLD and Ub(n)2-FOLD were photo-converted with UV light for 1 h on ice. The constructs for Ub(n)-NTAIL (NTAIL-Ub-G76V-Dendra) and Ub(n)-CTAIL (Ub-G76V-Dendra-CTAIL) were purified similarly without cleaving the His tag.

Ubx2 was expressed in yeast cells and purified from the membrane fraction after solubilization with 1% (w/v) decylmaltose neopentylglycol (DMNG) (Anatrace) by Ni-NTA and size-exclusion chromatography in 20 mM HEPES, 300 mM KCl, 5 mM magnesium acetate, 120 μM DMNG, and 0.5 mM TCEP, as described in (Stein et al., 2014). In all assays employing Ubx2, the protein was diluted to give a final concentration of 2 μM DMNG and 5 mM additional KCl. FLAG-26S was expressed in yeast cells and purified by FLAG antibody M2 agarose resin (Sigma) and size-exclusion chromatography, as described (de la Pena et al., 2018). Protein A-tagged 20S proteasomes were expressed in yeast cells and purified by IgG Sepharose 6 Fast Flow affinity resin (Cytiva). After removal of the tag, the protein was purified by ion-exchange and size-exclusion chromatography, as described (Leggett et al., 2005). His14-Uba1, His14-Ubr1, Ubc2, *Mus musculus* Ube1, the gp78^RING^-Ube2g2 fusion, and His14-SUMO-hUb were expressed and purified as described in (Ji *et al*., 2022). *S. cerevisiae* ubiquitin was purchased from Boston Biochem.

#### Dye labeling of proteins

The purified constructs were reduced with 10 mM TCEP and then incubated at pH 8.0 with an equal molar concentration (for Dendra constructs) or a 3-fold molar excess (NeD-sfGFP) of maleimide-conjugated DyLight dyes (Thermo Fisher). The reactions were kept in the dark at room temperature for 2 h before quenching with 20 mM dithiothreitol (DTT). The unreacted free dyes were removed with Dye Removal columns (Thermo Fisher, #22858).

#### Photoconversion of Ub-G76V-Dendra and Ub-G76V-Ub-G76V-Dendra

Purified Ub-G76V-Dendra Ub-G76V-Ub-G76V-Dendra containing fluorescent dye (∼4 mg/ml) was placed in a 200-µl PCR tube in an ice bath. A long-wavelength UV flashlight (395-410 nm, DULEX) was positioned 5 cm above the tube, and the sample was irradiated for 1 h, with occasional mixing. After photoconversion, Ub-G76V-Dendra was purified by gel filtration.

#### Ubiquitination of proteins

Ubiquitination of NeD-sfGFP was performed as described in (Ji *et al*., 2022), yielding (Ub)n-TAIL. Polyubiquitinated Dendra [(Ub)n-FOLD, (Ub)n-REFOLD, (Ub)n-NTAIL, (Ub)n-CTAIL, or (Ub)n2-FOLD] and free polyubiquitin chains [Ub(n)] were generated as described (Ji *et al*., 2022). Briefly, 10 µM Ub-G76V-Dendra was incubated with 1 µM mouse Ube1, 20 µM gp78^RING^-Ube2g2, and 500 µM purified human ubiquitin in 20 mM HEPES, pH 7.4, 100 mM NaCl, 2 mM DTT, 10 mM ATP, and 10 mM MgCl_2_, and incubated at 37°C. Ubiquitin and ATP were added in small aliquots every 30 min over the first 5 h. The reaction was then kept at 37°C overnight. After ubiquitination, the sample was concentrated and loaded onto a Superdex 200 Increase column equilibrated with SEC buffer. Fractions containing protein with 10-25 ubiquitin molecules were identified by SDS-PAGE analysis, pooled, and concentrated. For the experiment in figure S3H, Ub(n)-FOLD(s) substrate containing chains of 5-12 ubiquitin molecules were pooled.

#### Substrate degradation assays

100 nM of Ub(n)-FOLD, Ub(n)-TAIL, Ub(n)-REFOLD, Ub(n)-NTAIL, Ub(n)-CTAIL, or Ub(n)2-FOLD was mixed with 200 nM 26S proteasomes, 200 nM UN, 200 nM Cdc48, and 200 nM selected UBA-UBL or UBA-UBX proteins in 26S reaction buffer (60 mM HEPES, pH 7.4, 20 mM NaCl, 20 mM KCl, 10 mM MgCl_2_, 2.5% glycerol, and 1 mM TCEP) and 0.5 mg/ml protease-free bovine serum albumin (BSA). An ATP regeneration mixture (5 mM ATP, 16 mM creatine phosphate, 0.03 mg/mL creatine kinase) was added and the reaction was incubated at 30°C for 1 h unless mentioned otherwise. The samples were subjected to SDS-PAGE, followed by fluorescence scanning on an Odyssey imager (LI-COR) and Coomassie-blue staining. For experiments employing Ub(n)-TAIL and Ddi1, Ddi1 carried a mutation (D220N) that prevents cleavage of this substrate (Yip *et al*., 2020). For the analogous experiments with Ub(n)-FOLD, wild-type Ddi1 was used.

#### Substrate unfolding assays

Substrate unfolding experiments were performed as previously described (Twomey *et al*., 2019). Briefly, 400 nM of polyubiquitinated, photo-converted Dendra was mixed with 400 nM UN and 400 nM Cdc48 in 50 mM HEPES pH 7.5, 150 mM NaCl, 10 mM MgCl_2_, 0.5 mM TCEP, and 0.5 mg/ml protease-free bovine serum albumin (BSA). After 10-min pre-incubation at 30°C, an ATP regeneration mixture was added (10 mM ATP, 20 mM creatine phosphate, 100 µg/ml creatine kinase), and the fluorescence (excitation at 540 nm; emission at 580 nm; gain from 80 to 100) was measured at 15-s intervals for 30 min, using a Synergy Neo2 Multi-mode reader (BioTek).

For data acquired with the Synergy Neo2 Multi-mode reader instrument, the relative fluorescence at time t was calculated as (fluorescence at t) / (fluorescence at t_0_). The calculated relative fluorescence was plotted against time using Prism software (GraphPad).

#### ATPase assays

ATPase activity experiments were performed as previously described (Bodnar and Rapoport, 2017b), with the EnzCheck Phosphate Assay Kit (Invitrogen). Briefly, 20 µM 2-amino-6-mercapto-7-methylpurine riboside (MESG), 1 U/mL purine nucleoside phosphorylase, 200 nM UN, 200 nM UBA-UBX proteins or their variants, and 2 mM ATP were mixed in 1x EnzCheck buffer. After 10-min pre-incubation at 30°C, 200 nM Cdc48 was added, and absorbance at 360 nm was measured at 15-s intervals for 30 min, using a Synergy Neo2 Multi-mode reader (BioTek).

For data acquired with the Synergy Neo2 Multi-mode reader instrument, the ATPase activity of each reaction was calculated by the slope of absorbance curve from 5 to 15 min where the curve is linear. The calculated ATPase activity was plotted using Prism software (GraphPad).

#### Pull-down experiments

For all pull-down experiments using FLAG-26S proteasomes, 0.4 µM FLAG-26S proteasomes were mixed with 0.2 µM polyubiquitinated substrate, and 0.2 µM Rad23/Dsk2/Ddi1, in 26S reaction buffer (60 mM HEPES, pH 7.4, 20 mM NaCl, 20 mM KCl, 10 mM MgCl_2_, 2.5% glycerol, and 1 mM TCEP) supplemented with 5 mM ATPγS or 2 mM ADP·BeF_x_. 20 µl of the protein mixture were incubated with 5 µl pre-equilibrated FLAG antibody M2 agarose beads (Sigma) at 4°C for 1 h. The beads were then washed three times with reaction buffer containing the appropriate nucleotide. Bead-bound proteins were eluted with 25 µl of reaction buffer supplemented with 0.2 mg/ml 3xFLAG peptide (Bimake). The eluted material was subjected to SDS-PAGE, followed by fluorescence scanning on an Odyssey imager (LI-COR) and Coomassie-blue staining. For experiments employing Ub(n)-TAIL and Ddi1, Ddi1 carried a mutation (D220N) that prevents cleavage of this substrate (Yip *et al*., 2020). For the analogous experiments with Ub(n)-FOLD, wild-type Ddi1 was used.

For pull-down experiments with Npl4-FLAG or SBP tagged Rad23/Dsk2/Ddi1/Shp1/Ubx2/Ubx5, 1 µM polyubiquitinated substrate was mixed with 1 µM Cdc48, 1 µM HA-Ufd1, 1 µM Npl4-FLAG, 1 µM SBP tagged protein, in binding buffer (50 mM Tris-HCl, pH 8, 150 mM NaCl, 10 mM MgCl_2_, and 1 mM DTT) supplemented with 5 mM ATPγS or 2 mM ADP·BeF_x_. 20 µl of the protein mixture were then incubated with 10 µl pre-equilibrated FLAG antibody M2 agarose beads (Sigma) or streptavidin agarose beads (Thermo Fisher) at 4°C for 1 h. The beads were then washed three times with binding buffer containing the appropriate nucleotide. Bead-bound proteins were eluted with 25 µl of binding buffer supplemented with 0.05% Tween-20 and either 0.2 mg/ml 3xFLAG peptide (Bimake) or 2 mM biotin (Sigma). The eluted samples were subjected to SDS-PAGE, followed by fluorescence scanning on an Odyssey imager (LI-COR) and Coomassie-blue staining.

For the two-step pull-down experiments in Figure 4A, 5 µl pre-equilibrated streptavidin agarose beads were first incubated with 0.2 µM Cdc48 with a Strep tag at the C-terminus (Cdc48-strep), 0.2 µM polyubiquitinated substrate, 0.2 µM UN, and 0.2 µM Ubx5 in 26S reaction buffer supplemented with 5 mM ATPγS or 2 mM ADP·BeF_x_ for 1 h at 4°C. The beads were washed three times with binding buffer containing the appropriate nucleotide. The beads were then incubated with 0.2 µM 26S proteasomes and 0.2 µM Rad23 in 26S reaction buffer supplemented with the appropriate nucleotide for 1 h at 4°C. The beads were washed again and bead-bound proteins were eluted with 25 µl of binding buffer supplemented with 2 mM biotin.

For the two-step pull-down experiments in Figure 4B, 5 µl pre-equilibrated FLAG antibody M2 agarose beads were first incubated with 0.2 µM FLAG-26S proteasomes, 0.2 µM polyubiquitinated substrate, and 0.2 µM Rad23 in 26S reaction buffer supplemented with 2 mM ADP·BeF_x_ for 1 h at 4°C. The beads were washed three times with binding buffer containing ADP·BeF_x_. The beads were then incubated with 1 µM polyubiquitin in 26S reaction buffer supplemented with ADP·BeF_x_ for different time at 4°C. The beads were washed again, and bead-bound proteins were eluted with 25 µl of binding buffer supplemented with 0.2 mg/ml 3xFLAG peptide.

For the two-step pull-down experiments in Figure 4C, 5 µl pre-equilibrated FLAG antibody M2 agarose beads were first incubated with 0.2 µM FLAG-26S proteasomes, 0.2 µM polyubiquitinated substrate, and 0.2 µM Rad23 in 26S reaction buffer supplemented with 5 mM ATPγS or 2 mM ADP·BeF_x_ for 1 h at 4°C. The beads were washed three times with binding buffer containing the appropriate nucleotide. The beads were then incubated with 0.2 µM Cdc48, 0.2 µM UN, and 0.2 µM Ubx5 in 26S reaction buffer supplemented with the appropriate nucleotide for 1 h at 4°C. The beads were washed again and bead-bound proteins were eluted with 25 µl of binding buffer supplemented with 0.2 mg/ml 3xFLAG peptide.

For the two-step unfolding and degradation assays in Figures 4D and 4E, 20 µl pre-equilibrated FLAG antibody M2 agarose beads were first incubated with 1 µM FLAG-26S proteasomes, 2 µM polyubiquitinated substrate, and 2 µM Rad23 in 26S reaction buffer supplemented with 5 mM ATP for 1 h at 4°C. The beads were washed three times with binding buffer containing 1 mM ATP, and bead-bound proteins were eluted with 80 µl of binding buffer supplemented with 0.2 mg/ml 3xFLAG peptide. The proteasome concentration in the eluate was determined by SDS-PAGE gel. Cdc48, UN, and Ubx5 were added at the same molar concentration together with ATP and an ATP regeneration mix. The loss of fluorescence was monitored over time, and samples were taken at different time points and subjected to SDS-PAGE, followed by fluorescence scanning on an Odyssey imager (LI-COR) and Coomassie-blue staining.

#### Yeast cell viability assay

Each strain was grown at 25°C to an OD of 0.6-0.8. 1 mL of the cells was centrifuged and resuspended in 1 mL of fresh medium. The resuspended cells were serially diluted and plated on SD-medium agar.

#### Immunoprecipitation of proteasomes and Cdc48-UN from yeast lysates

Immunoprecipitation (IP) from lysates of *S. cerevisiae* cells was performed as described in (Tsuchiya et al., 2017) with some modifications. Briefly, a cell pellet corresponding to 40 OD_600_ was resuspended in 800 µl of lysis buffer (50 mM Tris-HCl, pH 7.5, 100 mM NaCl, 10% glycerol, 10 µM bortezomib, 10 mM iodoacetamide, 1x Protease Inhibitor Cocktail (Roche)) and mixed with 1 ml of acid-washed glass beads. The cells were lysed using a Mini-BeadBeater 96 (BIOSPEC) at 2,400 rpm for 4 min. The cell lysate was then spun at 860 g for 2 min at 4°C to remove unbroken cells and debris. The supernatant was supplemented with 1% TX-100 and incubated on ice for 30 min, before spinning at 20,000 g for 20 min. A small aliquot of the cleared lysate was used for SDS-PAGE and immunoblots with ubiquitin antibodies and SBP antibodies. The protein concentration in the cleared lysate was determined with a Protein Assay Dye Reagent (BioRad). Lysate containing ∼1 mg of proteins was incubated with 20 µl FLAG antibody agarose resin (Sigma, M2) or Pierce Anti-HA Magnetic Beads (Thermo Fisher) that was pre-equilibrated with lysis buffer containing 1% TX-100. After 1 h incubation at 4°C, the beads were washed three times with 1 ml of the lysis buffer supplemented with 1% TX-100. Bound material was eluted with 4 bead-volumes of elution buffer (50 mM HEPES, pH 7.4, 150 mM NaCl, 5 mM MgCl_2_, 0.5 mM TCEP, 0.1% TX-100) supplemented with 0.2 mg/ml 3xFLAG peptide or 1 mg/ml HA peptide. 20 µl of the total eluate was examined by SDS-PAGE and immunoblotting using ubiquitin antibodies, and FLAG antibodies or HA antibodies. The other 60 µl of the eluate were precipitated by adding 15 µl of 100% trichloroacetic acid (TCA) (Sigma). After 30-min incubation on ice, the samples were centrifuged at 14,000 rpm for 15 min at 4°C. The pellet was sequentially washed with 1 ml of ice-cold acetone and 1 ml of cold methanol, and processed for mass spectrometry.

#### Mass spectrometry

All samples were resuspended in 100µL of 100 mM 4-(2-Hydroxyethyl)-1-piperazinepropanesulfonic acid, 4-(2-Hydroxyethyl)piperazine-1-propanesulfonic acid, N-(2-Hydroxyethyl)piperazine-N′-(3-propanesulfonic acid) (EPPS), pH 8.5 and digested at 37°C with trypsin at a 100:1 protein-to-protease ratio overnight. The samples were desalted via StageTip, dried by vacuum centrifugation, and disolved in 5% acetonitrile, 5% formic acid for LC-MS/MS processing.

Mass spectrometry data were collected using a Exploris 480 mass spectrometer (Thermo Fisher Scientific, San Jose, CA) coupled with a Proxeon 1200 Liquid Chromatograph (Thermo Fisher Scientific). Peptides were separated on a 100 μm inner diameter microcapillary column packed with ∼25 cm of Accucore C18 resin (2.6 μm, 150 Å, Thermo Fisher Scientific). ∼1 μg was loaded onto the column. Peptides were separated using a 90min gradient of 4 to 30% acetonitrile in 0.125% formic acid with a flow rate of 520 nL/min. The scan sequence began with an Orbitrap MS1 spectrum with the following parameters: resolution 60,000, scan range 350−1350 Th, automatic gain control (AGC) target “standard”, maximum injection time “auto”, RF lens setting 40%, and centroid spectrum data type. The top twenty precursors were selected for MS2 analysis which consisted of HCD high-energy collision dissociation with the following parameters: resolution 15,000, AGC was set at “standard”, maximum injection time “auto”, isolation window 0.7 Th, normalized collision energy (NCE) 28, and centroid spectrum data type. In addition, unassigned and singly charged species were excluded from MS2 analysis and dynamic exclusion was set to 90 s.

Mass spectra were processed using a Comet-based in-house software pipeline. MS spectra were converted to mzXML using a modified version of ReAdW.exe. Database searching with *S. cerevisiae* UniProt database were performed as described above. PSM filtering was performed using a linear discriminant analysis, as described previously (Huttlin et al., 2010), while considering the following parameters: XCorr, ΔCn, missed cleavages, peptide length, charge state, and precursor mass accuracy. Peptide-spectral matches were identified, quantified, and collapsed to a 5% FDR and then further collapsed to a final protein-level FDR of 5%. Protein assembly was also guided by principles of parsimony to produce the smallest set of proteins necessary to account for all observed peptides.

For each sample, the spectral counts were normalized to the first sample based on the average ratio of proteasome subunits spectral counts (for proteasome pull-down) or the average ratio of Cdc48, Ufd1, and Npl4 spectral counts (for Cdc48-UN pull-down). For each protein detected, the ratio of its counts in cdc48-3 versus wt, cdc48-3 ΔUbx5 versus cdc48-3, cdc48-3 + Ubx5 versus cdc48-3, and cdc48-3 ΔRad23 ΔDsk2 versus cdc48-3 were calculated. The P values were then plotted against these ratios.

### QUANTIFICATION AND STATISTICAL ANALYSIS

Quantification of bands after fluorescence scanning of gels was carried out using the ImageStudio software (LI-COR). For each lane, a rectangular box was selected to determine the total intensity of a band. In the case of polyubiquinated species, the box included the smear corresponding to the entire range of ubiquitin molecules in the chains. The box size was kept constant for all bands on the same gel. An additional box of the same size was drawn over an empty region to determine background intensity. Signal intensity of each band was calculated as (total intensity –background intensity). The resulting signal intensity was normalized to a designated lane to calculate the relative signal intensity.

The percentage of substrate degraded over time was quantified by determining the fluorescence in peptides after subtracting background fluorescence. These numbers were compared to the total fluorescence in each lane of the SDS gel. It should be noted that, in each experiment, the total florescence in different lanes was constant, with differences smaller than 20%.

Band intensities on Coomassie blue-stained gels were quantified using ImageJ (NIH). Background subtraction and normalization were performed as described for gels analyzed by fluorescence scanning.

