## Supplement material for "Bidirectional substrate shuttling between the 26S proteasome and the Cdc48 ATPase promotes protein degradation"

### Supplemental Material

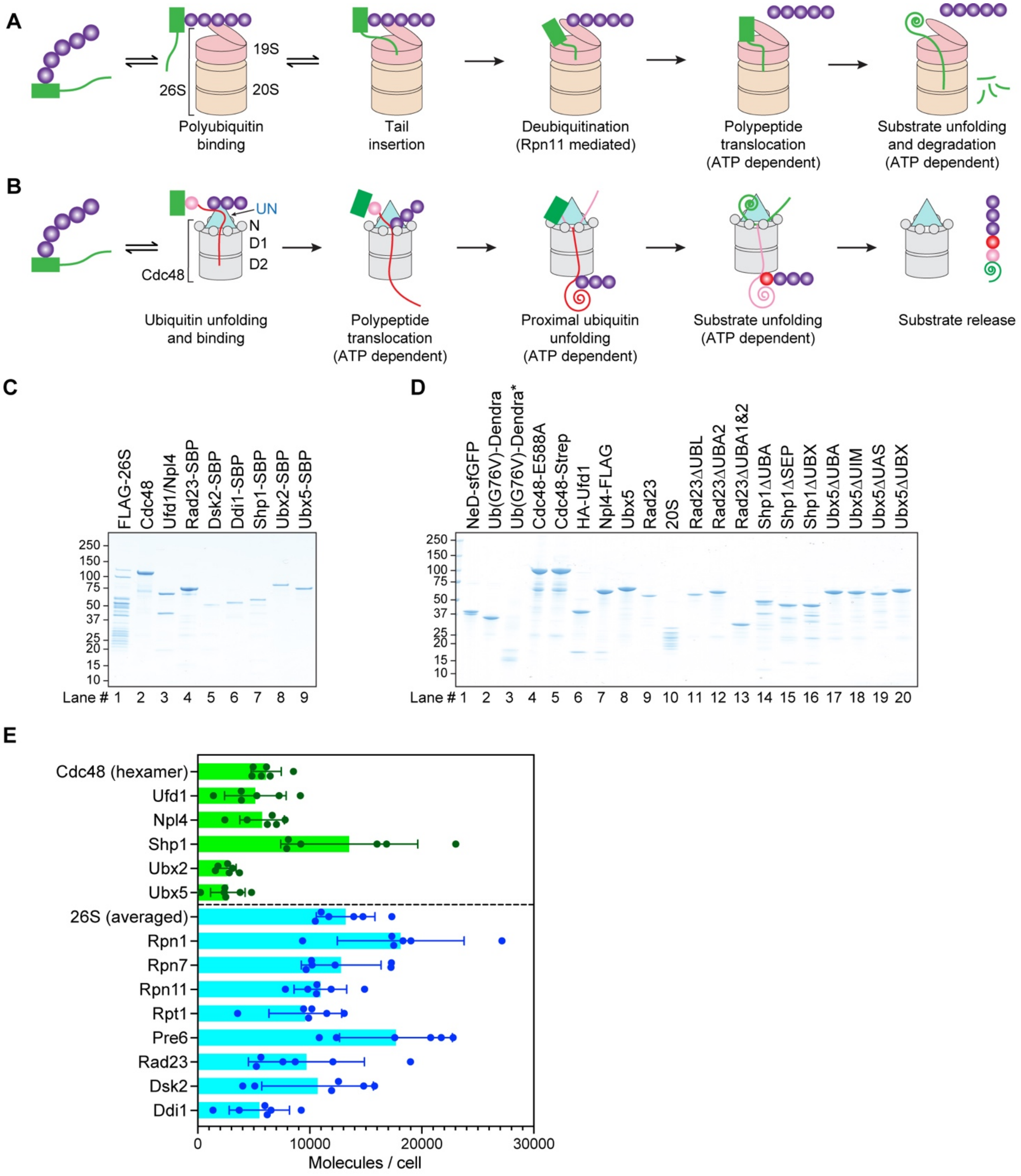


**Figure S1 (related to Figures 1-5). Mechanisms of the 26S proteasome and Cdc48 ATPase complex and purity and abundance of cofactors used in this study.**

(**A**) Scheme of protein degradation by the proteasome. A polyubiquitinated substrate with a flexible segment binds to the 19S regulatory subunit. The segment inserts into the ATPase ring, the ubiquitin chain is removed by the DUB Rpn11, and the substrate is translocated into the 20S proteolytic chamber and degraded into peptides.

(**B**) Scheme of the function of the Cdc48-UN ATPase complex. A folded, polyubiquitinated substrate binds to the ATPase complex through the Ufd1/Npl4 (UN) cofactor. One of the ubiquitin molecules (initiator, in red) is unfolded and projects its N-terminus across both ATPase rings. ATP hydrolysis by the D2 ring ATPases moves the ubiquitin molecules linked to the C-terminus of the initiator (proximal ubiquitin molecules, in pink) and ultimately the substrate through the central pore and causes their unfolding.

(**C**) The indicated proteins were purified and subjected to SDS-PAGE followed by staining with Coomassie-blue.

(**D**) as in (C), but with additional proteins.

(**E**) Protein abundance in *S. cerevisiae* cells, based on quantitative mass spectrometry performed in six different studies (de Godoy et al., 2008; Kulak et al., 2014; Nagaraj et al., 2012; Peng et al., 2012; Thakur et al., 2011; Webb et al., 2013). For the 26S proteasome, the abundance of selected constituents is shown, as well as an average based on all constituents.

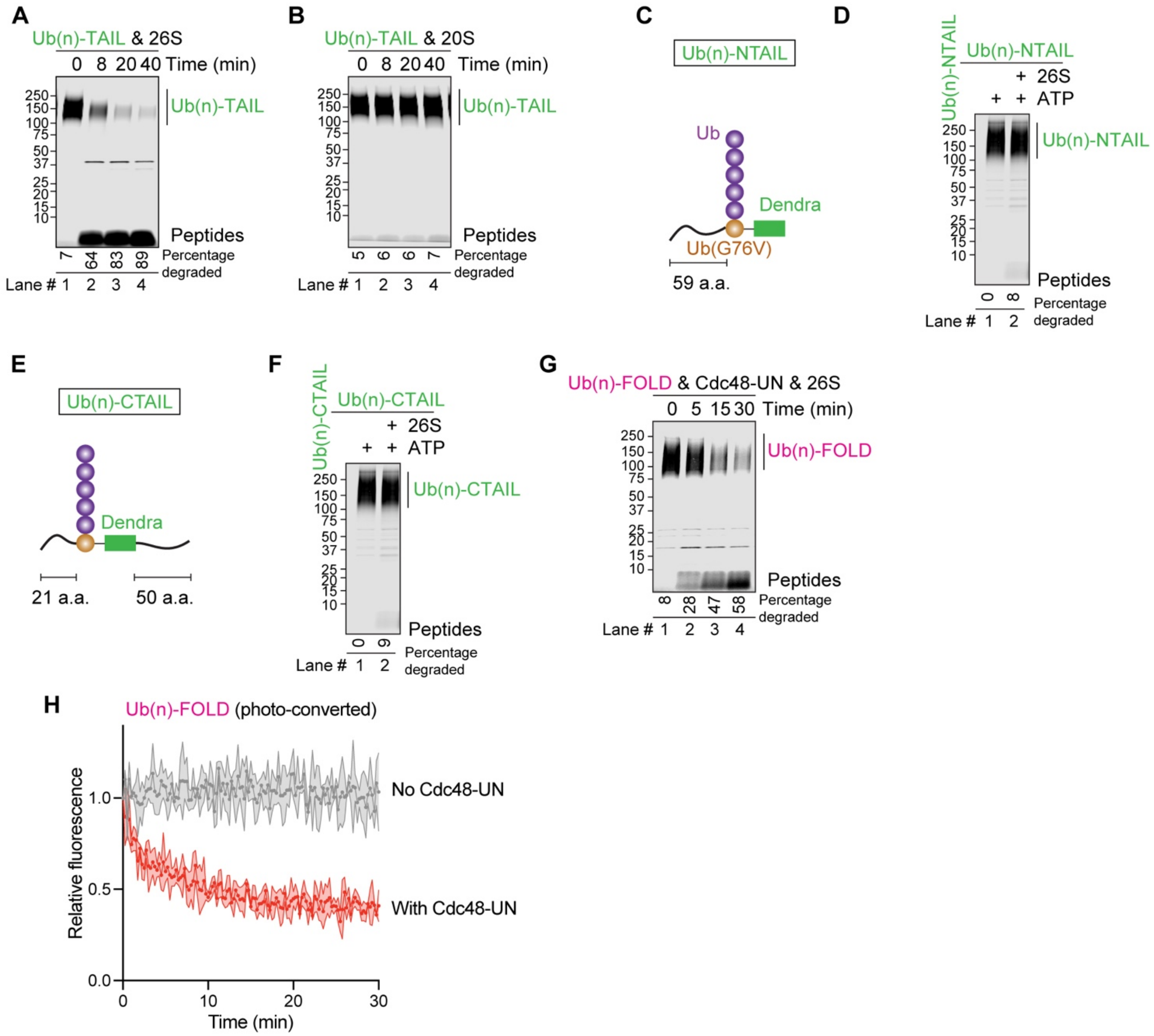


**Figure S2 (related to Figure 1). Proteasomal degradation of model substrates.**

(**A**) Time course of degradation of Dylight800-labeled Ub(n)-TAIL by the 26S proteasome. The samples were analyzed by SDS-PAGE, followed by fluorescence scanning. The percentage of substrate degraded was quantified by determining the fluorescence in peptides (after background subtraction) and comparing it to the total fluorescence in each lane of the SDS gel.

(**B**) As in (A), but for the 20S core particle, i.e. in the absence of the 19S regulatory subunit.

(**C**) Scheme of the model substrate Ub(n)-NTAIL. The ubiquitin (Ub) mutant Ub-G76V with a flexible segment of 59 amino acids at the N-terminus was fused to Dendra with an 8 amino acid linker. A ubiquitin chain was attached to Ub-G76V.

(**D**) Ub(n)-NTAIL, labeled with the fluorescent dye Dylight 800, was incubated with 26S proteasomes for 60 min and analyzed by SDS-PAGE and fluorescence scanning. The percentage of substrate degraded was quantified as in (A).

(**E**) Scheme of the model substrate Ub(n)-CTAIL. The ubiquitin (Ub) mutant Ub-G76V was fused to Dendra with an 8 amino acid linker. The construct contains a flexible segment of of 50 amino acids at the C-terminus. A ubiquitin chain was attached to Ub-G76V.

(**F**) As in (D), but with Dylight 800-labeled Ub(n)-CTAIL.

(**G**) As in (A), but with photo-converted Ub(n)-FOLD in the presence of 26S proteasomes and Cdc48-UN.

(**H**) Ub(n)-FOLD, containing photo-cleaved Dendra, was incubated with or without Cdc48-UN complex. The loss of fluorescence was followed over time.

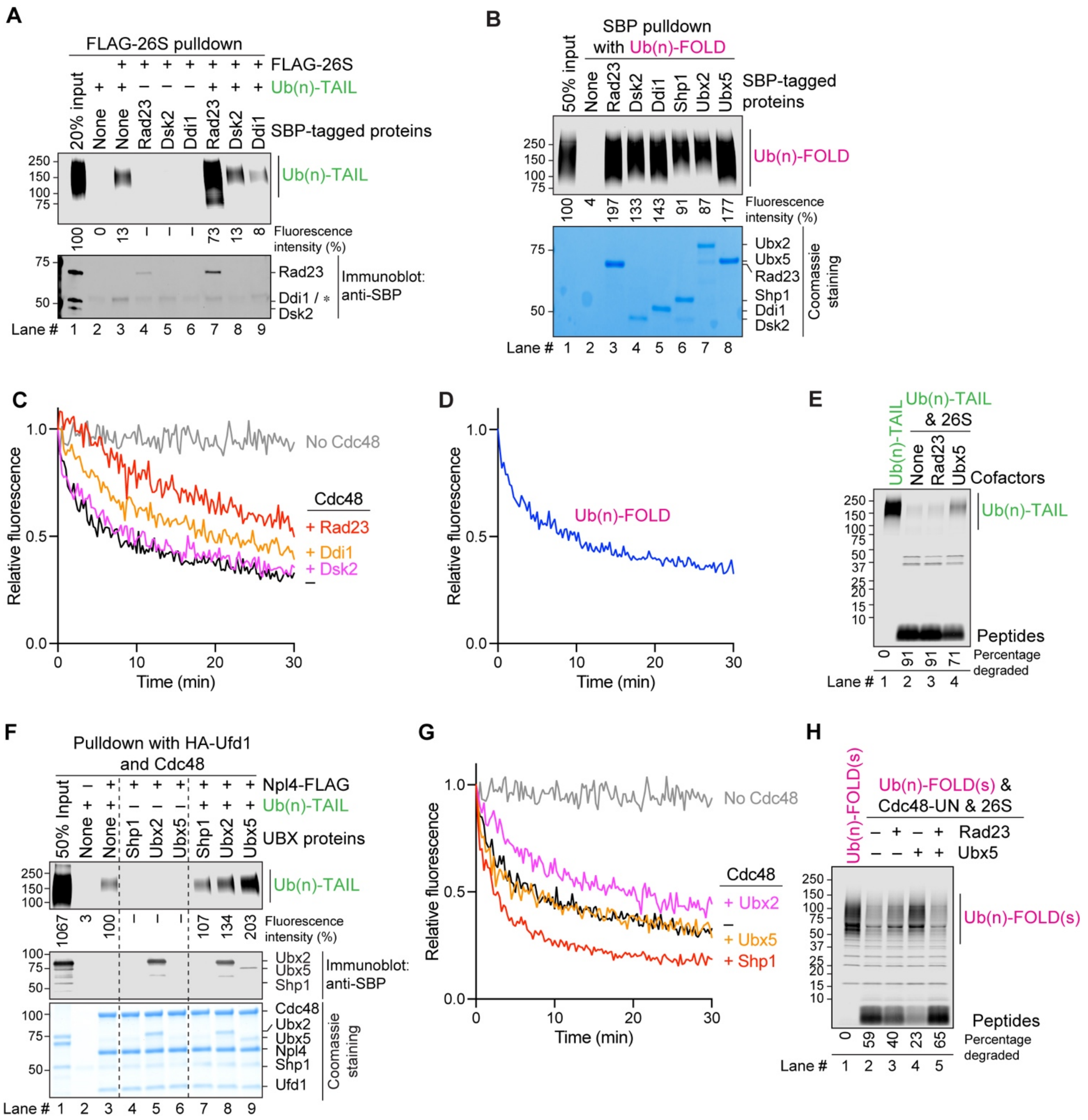


**Figure S3 (related to Figures 2, 3, and 5). Testing proteasome and Cdc48 cofactors for substrate binding, unfolding, and degradation of model substrates.**

(**A**) Testing UBA-UBL proteins in the recruitment of Ub(n)-FOLD to the proteasome. FLAG-tagged 26S proteasomes were incubated with fluorescently labeled Ub(n)-FOLD and SBP-tagged versions of Rad23, Dsk2, or Ddi1, as indicated. After immunoprecipitation with FLAG antibodies, the samples were analyzed by SDS-PAGE followed by fluorescence scanning and blotting for SBP. The numbers below the gel give the intensity of the fluorescent bands relative to 20% input.

(**B**) Dylight800-labeled Ub(n)-FOLD was incubated with SBP-tagged cofactors of the 26S proteasome or Cdc48. The cofactors were retrieved with streptavidin beads, and the bound material analyzed by SDS-PAGE, followed by fluorescence scanning and Coomassie-blue staining. Bound fluorescent substrate was quantified relative to 50% input.

(**C**) Photo-converted Ub(n)-FOLD was incubated with Cdc48 ATPase, UN, and cofactors of the proteasome (Rad23, Dsk2, Ddi1). Where indicated, the cofactors or Cdc48 were omitted. The loss of fluorescence caused by unfolding of Dendra was followed over time.

(**D**) Photo-converted Ub(n)-FOLD was first unfolded by Cdc48-UN, as shown by the loss of fluorescence. The unfolded protein was then tested for degradation (**Figure 2E**).

(**E**) Dylight800-labeled Ub(n)-TAIL was incubated with 26S proteasomes alone, or together with Rad23 or Ubx5. After 60 min, the samples were analyzed by SDS-PAGE, followed by fluorescence scanning. The percentage of substrate degraded was quantified by determining the fluorescence in peptides (after background subtraction) and comparing it to the total fluorescence in each lane of the SDS gel.

(**F**) Fluorescently labeled Ub(n)-TAIL was incubated with Cdc48, Ufd1, and FLAG-tagged Npl4. Where indicated, SBP-tagged versions of Shp1, Ubx2, or Ubx5 were added. The Cdc48 complex was retrieved with beads containing FLAG antibodies, and the bound material analyzed by SDS-PAGE, followed by fluorescence scanning, blotting for SBP, and staining with Coomassie-blue. Bound fluorescent substrate was quantified relative to the sample lacking cofactors (lane 3) (numbers below the gel).

(**G**) As in (C), but with Cdc48 cofactors Shp1, Ubx2, and Ubx5.

(**H**) Fluorescently labeled, photo-converted Ub(n)-FOLD(s) carrying 5-12 ubiquitins was incubated with 26S proteasomes, Cdc48-UN, Rad23, and Ubx5 as indicated. After 60 min, the samples were analyzed by SDS-PAGE and fluorescence scanning. Quantification was done as in (E).

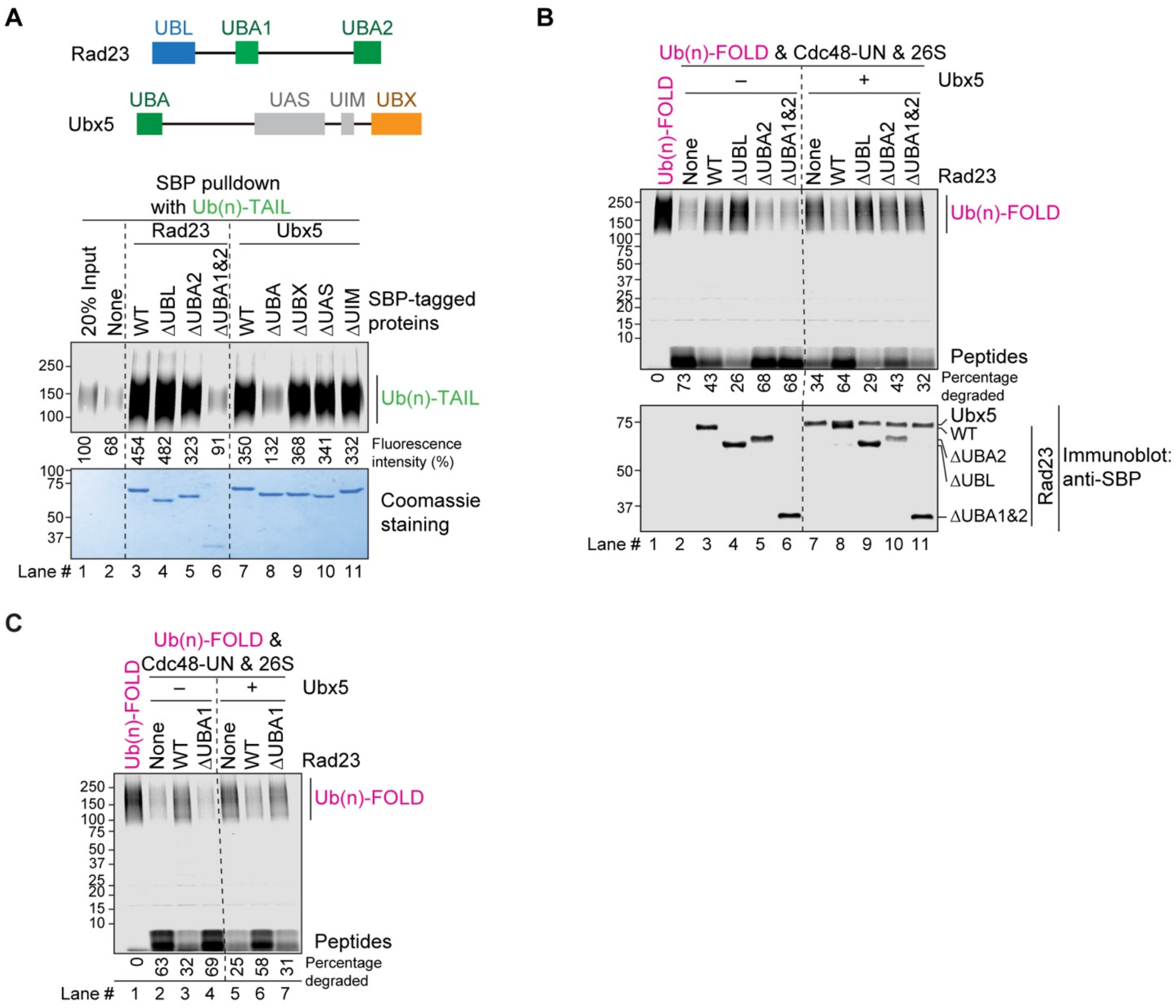


**Figure S4 (related to Figure 4). Domains of Rad23 and Ubx5 required for substrate binding and degradation.**

(**A**) The scheme shows the domains of Rad23 and Ubx5. In the experiment below, Dylight 800-labeled Ub(n)-TAIL was incubated with SBP-tagged wild-type (WT) Rad23 or Ubx5, or with the indicated deletion mutants. Rad23ΔUBA2 lacks the second UBA domain and ΔUBA1&2 lacks both UBA domains. Material bound to streptavidin beads was analyzed by SDS-PAGE, followed by fluorescence scanning and Coomassie-blue staining. Bound fluorescent substrate was quantified relative to 20% input (numbers below the gel).

(**B**) The degradation of Ub(n)-FOLD was tested with 26S proteasomes, the Cdc48-UN complex, and either wild-type Rad23 or the indicated deletion mutants. Where indicated, Ubx5 was also present. The percentage of substrate degraded was quantified by determining the fluorescence in peptides (after background subtraction) and comparing it to the total fluorescence in each lane of the SDS gel.

(**C**) As in (B), but with Rad23ΔUBA1 which lacks the first UBA domain.

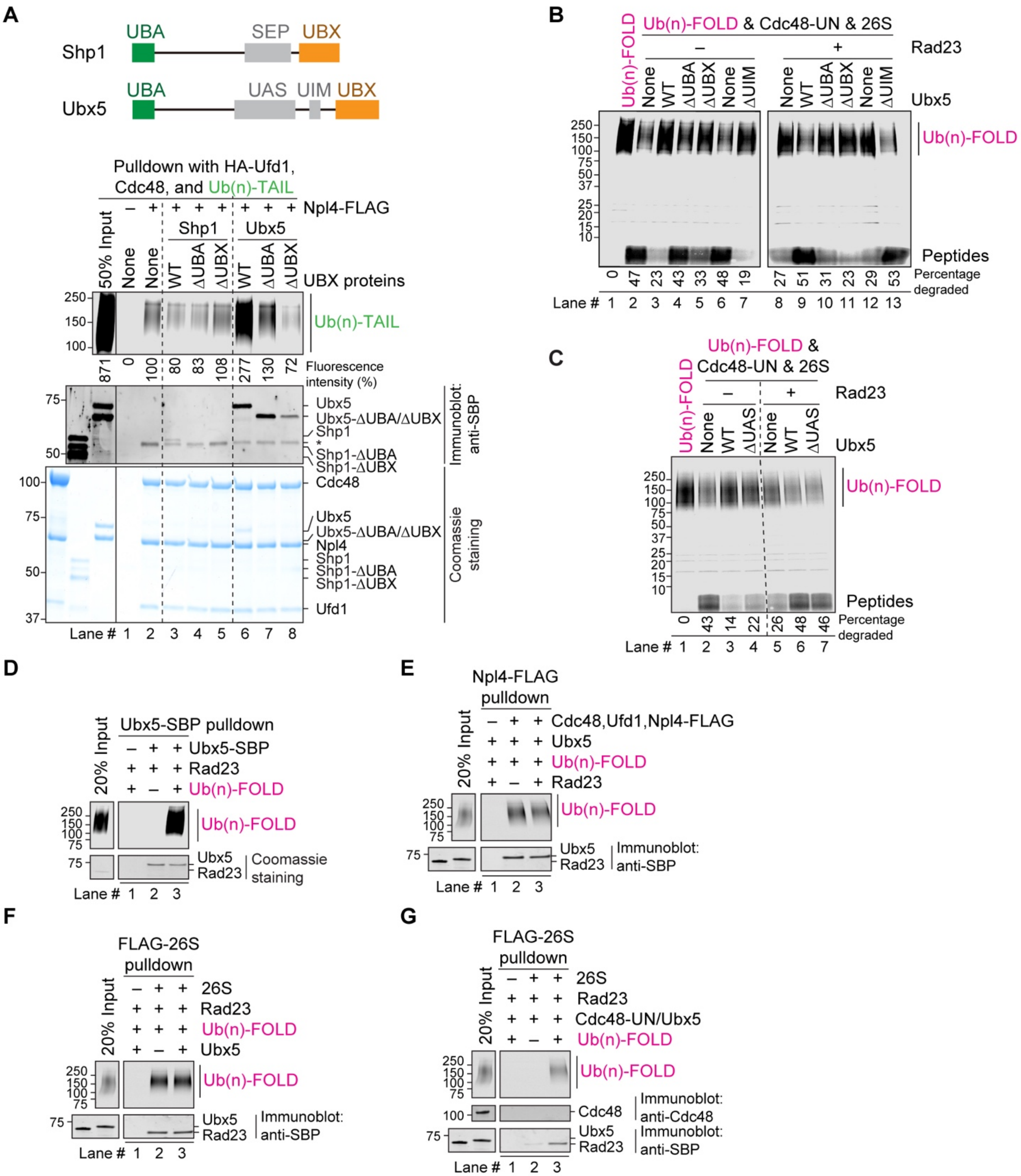


**Figure S5 (related to Figure 4). Interaction of cofactors with the proteasome and Cdc48 ATPase complex.**

(**A**) The scheme shows the domains of Shp1 and Ubx5. In the experiment below, Dylight800-labeled Ub(n)-TAIL was incubated with Cdc48, Ufd1, and FLAG-tagged Npl4, as indicated. SBP-tagged versions of wild-type (WT) Shp1 or Ubx5, or of deletion mutants, were added. The Cdc48 complex was retrieved with beads containing FLAG antibodies, and the bound material analyzed by SDS-PAGE, followed by fluorescence scanning, staining with Coomassie-blue, and blotting for SBP. Bound fluorescent substrate was quantified relative to the sample lacking cofactors (lane 2) (numbers below the gel). Note that substrate and Ubx5 were most efficiently bound when present together. The star in the anti-SBP blot indicates a non-specific band.

(**B**) The degradation of Ub(n)-FOLD was tested with 26S proteasomes, the Cdc48-UN complex, and either wild-type Ubx5 or the indicated deletion mutants. Where indicated, Rad23 was also present. The percentage of substrate degraded was quantified by determining the fluorescence in peptides (after background subtraction) and comparing it to the total fluorescence in each lane of the SDS gel.

(**C**) As in (B), but with an Ubx5 mutant lacking the UAS domain (Ubx5ΔUAS).

(**D**) Ubx5-SBP was incubated with Rad23 with or without Ub(n)-FOLD and retrieved with streptavidin beads. The bound material was analyzed by SDS-PAGE, followed by fluorescence scanning and staining with Coomassie-blue.

(**E**) Dylight800-labeled Ub(n)-FOLD was incubated with Cdc48, Ufd1, FLAG-tagged Npl4, and Ubx5-SBP. SBP-tagged Rad23 was added as indicated. The Cdc48 complex was retrieved with beads containing FLAG antibodies, and the bound material analyzed by SDS-PAGE, followed by fluorescence scanning, and blotting for SBP.

(**F**) Dylight800-labeled Ub(n)-FOLD was incubated with FLAG-26S proteasomes and Rad23-SBP. SBP-tagged Ubx5 was added as indicated. The proteasomes were retrieved with beads containing FLAG antibodies, and the bound material analyzed by SDS-PAGE, followed by fluorescence scanning, and blotting for SBP.

(**G**) FLAG-26S proteasomes and Rad23-SBP were incubated with Cdc48-UN and Ubx5-SBP. Dylight800-labeled Ub(n)-FOLD were added as indicated. The proteasomes were retrieved with beads containing FLAG antibodies, and the bound material analyzed by SDS-PAGE, followed by fluorescence scanning, and blotting for SBP and Cdc48.

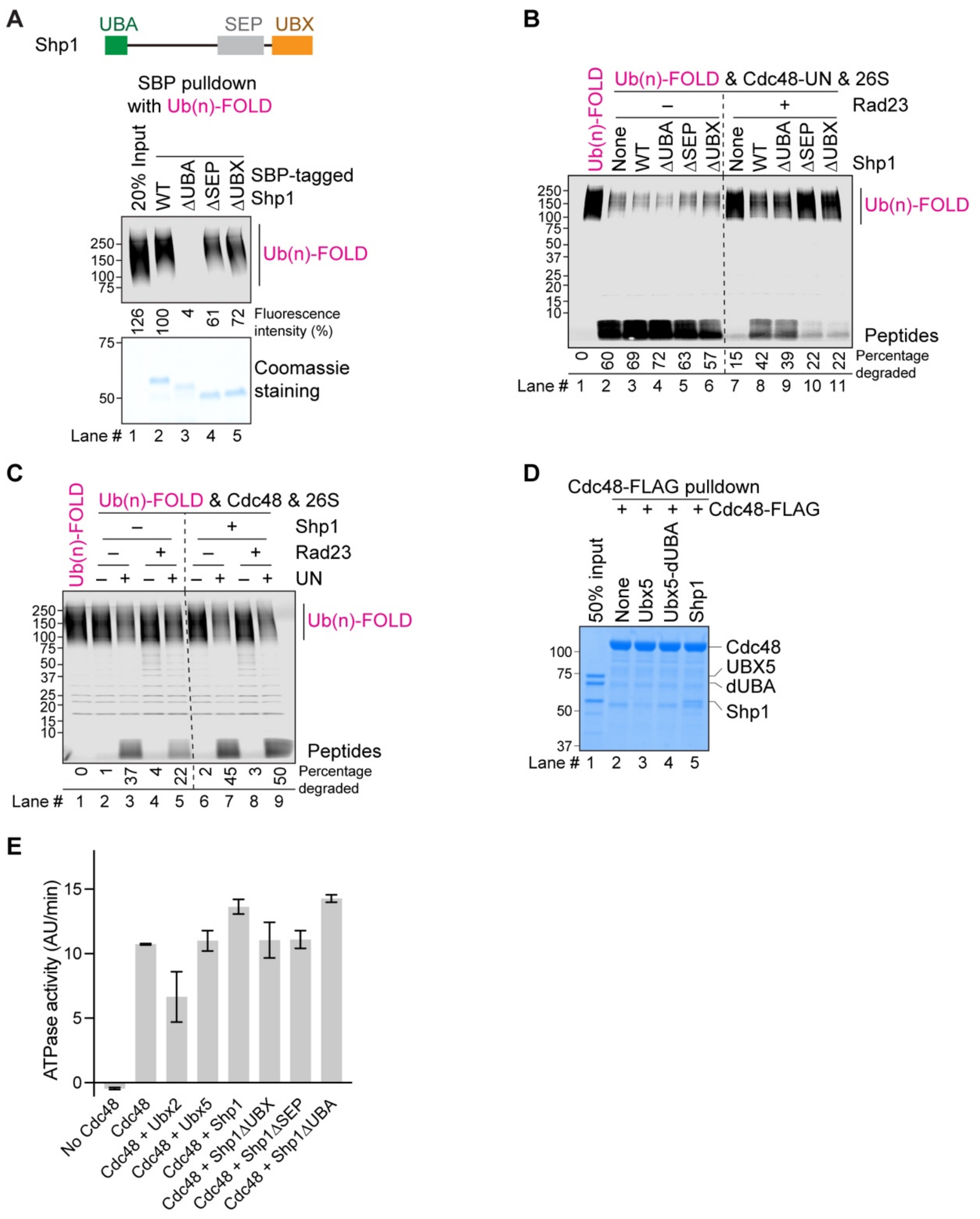


**Figure S6 (related to Figure 5). Testing the Cdc48-cofactor Shp1 in substrate binding and degradation.**

(**A**) The scheme shows the domains of Shp1. In the experiment below, Ub(n)-FOLD was incubated with SBP-tagged wild-type Shp1 or deletion mutants. Streptavidin-bound material was analyzed by SDS-PAGE, followed by fluorescence scanning and Coomassie-blue staining.

Bound fluorescent substrate was quantified relative to the sample containing wild-type (WT) Shp1 (lane 2) (numbers below the gel).

(**B**) Fluorescently labeled Ub(n)-FOLD was incubated with 26S proteasomes, Cdc48-UN, Rad23, and either WT Shp1 or deletion mutants, as indicated. After 60 min, the samples were analyzed by SDS-PAGE, followed by fluorescence scanning. Note that only the deletion of the SEP or UBX domains affects the activity of Shp1. The percentage of substrate degraded was quantified by determining the fluorescence in peptides (after background subtraction) and comparing it to the total fluorescence in each lane of the SDS gel.

(**C**) The degradation of Ub(n)-FOLD was tested in the presence of the indicated components. Note that the function of Shp1 depends on the presence of UN. Quantification was done as in (B).

(**D**) Cdc48-FLAG was incubated with Ubx5, Ubx5ΔUBA, or Shp1, as indicated. The Cdc48 complex was retrieved with beads containing FLAG antibodies, and the bound material analyzed by SDS-PAGE, followed by staining with Coomassie-blue.

(**E**) The ATPase activity of Cdc48 was measured in the presence of UN and the indicated proteins. Shown are means and standard errors of three experiments.

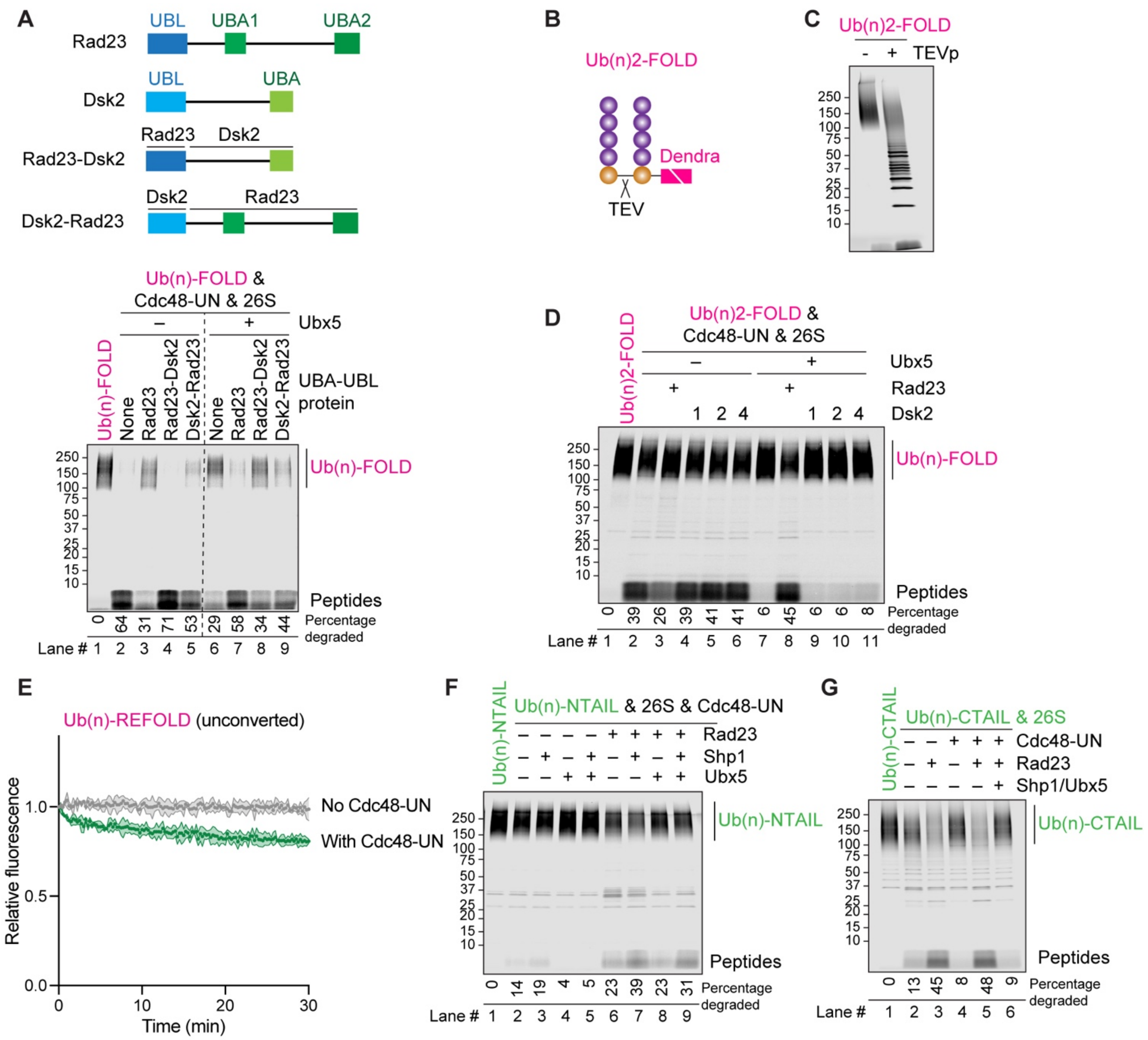


**Figure S7 (related to Figure 5). Testing the degradation of model substrates.**

(**A**) The scheme shows the domain organization of Rad23, Dsk2, and of chimeras of the two proteins. Dylight800-labeled Ub(n)-FOLD was incubated with 26S proteasomes, Cdc48-UN, Ubx5, and either wild-type Rad23 or chimeras in which the UBL domain of Dsk2 was replaced with that of Rad23 (Rad23-Dsk2) or the Dsk2 segment following the UBL domain was replaced by the corresponding segment of Rad23 (Dsk2-Rad23) (see scheme). After 60 min, the samples were analyzed by SDS-PAGE, followed by fluorescence scanning. The percentage of substrate degraded was quantified by determining the fluorescence in peptides (after background subtraction) and comparing it to the total fluorescence in each lane of the SDS gel.

(**B**) Scheme of the model substrate Ub(n)2-FOLD. Two molecules of the ubiquitin (Ub) mutant Ub-G76V, with a TEV protease cleavage site in between, were fused with an 8 amino acid linker to Dendra. Dendra was photo-cleaved into two polypeptide fragments (indicated by a white line). Each Ub-G76V molecule carried a ubiquitin chain.

(**C**) Dylight800-labeled Ub(n)2-FOLD was incubated with a 10-fold excess of TEV protease at room temperature. After 60 min, the samples were analyzed by SDS-PAGE, followed by fluorescence scanning. Note that the molecular weight of the substrate decreased, consistent with the removal of a ubiquitin chain by the TEV protease.

(**D**) Fluorescently labeled Ub(n)2-FOLD was incubated with 26S proteasomes, Cdc48-UN, Ubx5, Rad23, and different molar ratios of Dsk2, as indicated. After 60 min, the samples were analyzed by SDS-PAGE, followed by fluorescence scanning. Quantification as in (A).

(**E**) Ub(n)-REFOLD, containing intact Dendra (unconverted), was incubated with or without Cdc48-UN complex. The loss of fluorescence was followed over time. Note that there is only a small change, as the protein rapidly refolds.

(**F**) Dylight800-labeled Ub(n)-NTAIL was incubated with 26S proteasomes, Cdc48-UN, Rad23, Shp1, and Ubx5 as indicated. After 60 min, the samples were analyzed by SDS-PAGE, followed by fluorescence scanning. Quantification as in (A).

(**G**) Dylight800-labeled Ub(n)-CTAIL was incubated with 26S proteasomes, Cdc48-UN, Rad23, Shp1, and Ubx5 as indicated. After 60 min, the samples were analyzed by SDS-PAGE, followed by fluorescence scanning. Quantification as in (A).
